# Integrated multi-model analysis of intestinal inflammation exposes key molecular features of preclinical and clinical IBD

**DOI:** 10.1101/2024.08.14.607902

**Authors:** Miguel González-Acera, Jay V. Patankar, Lena Erkert, Roodline Cineus, Reyes Gámez-Belmonte, Tamara Leupold, Marvin Bubeck, Li-Li Bao, Martin Dinkel, Ru Wang, Heidi Limberger, Iris Stolzer, Katharina Gerlach, Fabrizio Mascia, Kristina Koop, Christina Plattner, Gregor Sturm, Benno Weigmann, Claudia Günther, Stefan Wirtz, Kai Hildner, Anja A. Kühl, Britta Siegmund, Raja Atreya, The IBDome Consortium, Ahmed N. Hegazy, Zlatko Trajanoski, Markus F. Neurath, Christoph Becker

**Author notes:** Corresponding author **Address correspondence to:** Christoph Becker, PhD Department of Medicine 1, Universitätsklinikum Erlangen Friedrich-Alexander-Universität Erlangen-Nürnberg (FAU) Erlangen, Germany. Share first-authorship (Miguel González-Acera, Jay V. Patankar).

## Abstract

**Background:** Inflammatory bowel disease (IBD) is a chronic inflammatory condition of the intestine with a complex and multifaceted pathogenesis. While various animal models exist to study specific disease mechanisms relevant to human IBD, a comprehensive comparative framework linking these to IBD pathophysiology is lacking.

**Objective:** In our study, we aimed at providing a framework that delineates common and unique features encountered in 13 widely used mouse models comparing them with human IBD to identify translatable pathways in model-cohort pairs. Another aim of our study was to provide an explorable resource for looking up gene and pathway level changes in mouse models assisting in hypothesis testing and minimizing animal burden abiding by the 3R principals.

**Design:** We employed comparative transcriptomic analyses with curated and *a priori* statistical correlative methods between mouse models versus established as well as own patient datasets at both bulk and single cell levels.

**Results:** We identify IBD-related pathways, ontologies, and cellular processes that are translatable between mouse models and patient cohorts. Moreover, we identify, known and novel IBD-associated subcellular mechanisms and how they are recapitulated in specific mouse models.

**Conclusion:** Our findings provide a valuable resource for selecting the most appropriate experimental paradigm to model unique features of IBD pathomechanisms, allowing analysis at the tissue, cellular, and subcellular levels.

**What is already known on this topic:** Preclinical modelling of IBD is key to the discovery of pathomechanisms and the evaluation of therapeutic approaches. However, individual models do not recapitulate the complexity of the disease and comprehensive studies comparing modelling paradigms with human IBD are lacking.

**What this study adds:** Our study provides a comparative analysis of thirteen commonly used intestinal inflammation models, identifying core-conserved pathways between mouse models and IBD patient cohorts. In addition, our study shows how specific pathways involved in IBD are recapitulated in specific mouse models and introduces a web tool to analyse the models.

**How this study might affect research, practice or policy:** By identifying conserved and discrepant pathways between specific mouse models and IBD patient cohorts, our analysis platform provides an invaluable resource for translational IBD research.

## Introduction

Inflammatory bowel disease (IBD) represents a group of debilitating and chronic gastrointestinal disorders that include Crohn’s disease (CD) and ulcerative colitis (UC). Despite significant progress in our understanding of IBD, the precise aetiology remains elusive. IBD is multifactorial and features a) an exaggerated and dysregulated immune response, b) barrier dysfunction and microbial dysbiosis, c) genetic susceptibility, and d) other unknown environmental causes [1, 2, 3]. Recent genome-wide association studies have identified over 200 IBD-associated genomic loci, some that directly confer increased susceptibility to IBD [4, 5]. However, causal relationships have only been established for a few genetic variants, and the contributions of other factors in triggering IBD has suffered from the “chicken or the egg” dilemma. This, together with the lack of shared molecular, prognostic, and etiological factors, has led investigators to hypothesize that IBD is a spectrum of diverse underlying disease mechanisms resulting in overlapping clinical presentation [6].

To unravel the mechanisms underlying the complex nature of IBD, various mouse models have been established. Modelling complex diseases in preclinical models is challenging, yet enables a simulation of disparate disease aspects in a controlled context. Experimental models of intestinal inflammation are based on three broad approaches: (I) disruption of the intestinal epithelial barrier, (II) activation of an aberrant immune response, and (III) alteration of microbial homeostasis. Each approach replicates specific components of disease aetiology, providing invaluable tools for exploring the IBD heterogeneity. However, none of these models fully reflect the complexity of human disease. This is exemplified by the distinct portrayal of particular pathobiological characteristics in each experimental model. Deciphering disease biology and the developing novel therapeutic strategies therefore relies on the choice of appropriate model to investigate the endophenotypes of concern.

Leveraging preclinical modelling of intestinal inflammation and exposing conserved IBD relevant molecular features, here we generated transcriptomic data from thirteen commonly used mouse models of intestinal inflammation. By employing both correlative statistics and curation-based measures of comparison across the regulomes of each model we uncovered shared and unique co-regulated modules related to ciliary processes, inflammatory response, carboxylic acid metabolism, mitochondriopathy, and synaptic processes. Comparative transcriptomics between mouse models and multiple public and own IBD cohorts identified age-specific signatures, suggesting the existence of conserved regulatory mechanisms that are differentially affected in paediatric versus adult patients. Our findings offer a framework for comparing and selecting appropriate modelling paradigms that align with conserved disease processes, paving the way for discovering novel translatable therapeutic targets.

## Results

### Modelling transcriptomes across experimental intestinal inflammation

Modelling of intestinal inflammation in mice can be broadly classified into three categories: barrier damage, immune modulation, and infection. To explore disease mechanisms, we recreated mouse models representing these categories on a C57BL/6J strain. We established cohorts for thirteen models (Fig. 1A): AcDSS and cDSS colitis [7], *Casp8*^ΔIEC^Ile and *Casp8*^ΔIEC^Col ileitis and colitis [8] for barrier damage; TC [9], OxC [7], AcTNBS and cTNBS [7], *Tnf*^ΔARE^Ile and *Tnf*^ΔARE^Col [10] for immune modulation; Everm [11], Hhepa [12], and Crode [13] for infection. An overview of the models and abbreviation details, are provided in Supplemental Table ST1. Models were pre-screened using colonoscopy, *in vivo* imaging, and faecal bacterial load measurement (Supplemental Fig. S1A, B). Post-euthanasia histology and transgene expression provided tissue-level insights (Supplemental Fig. S1A, B).

**Figure 1.**
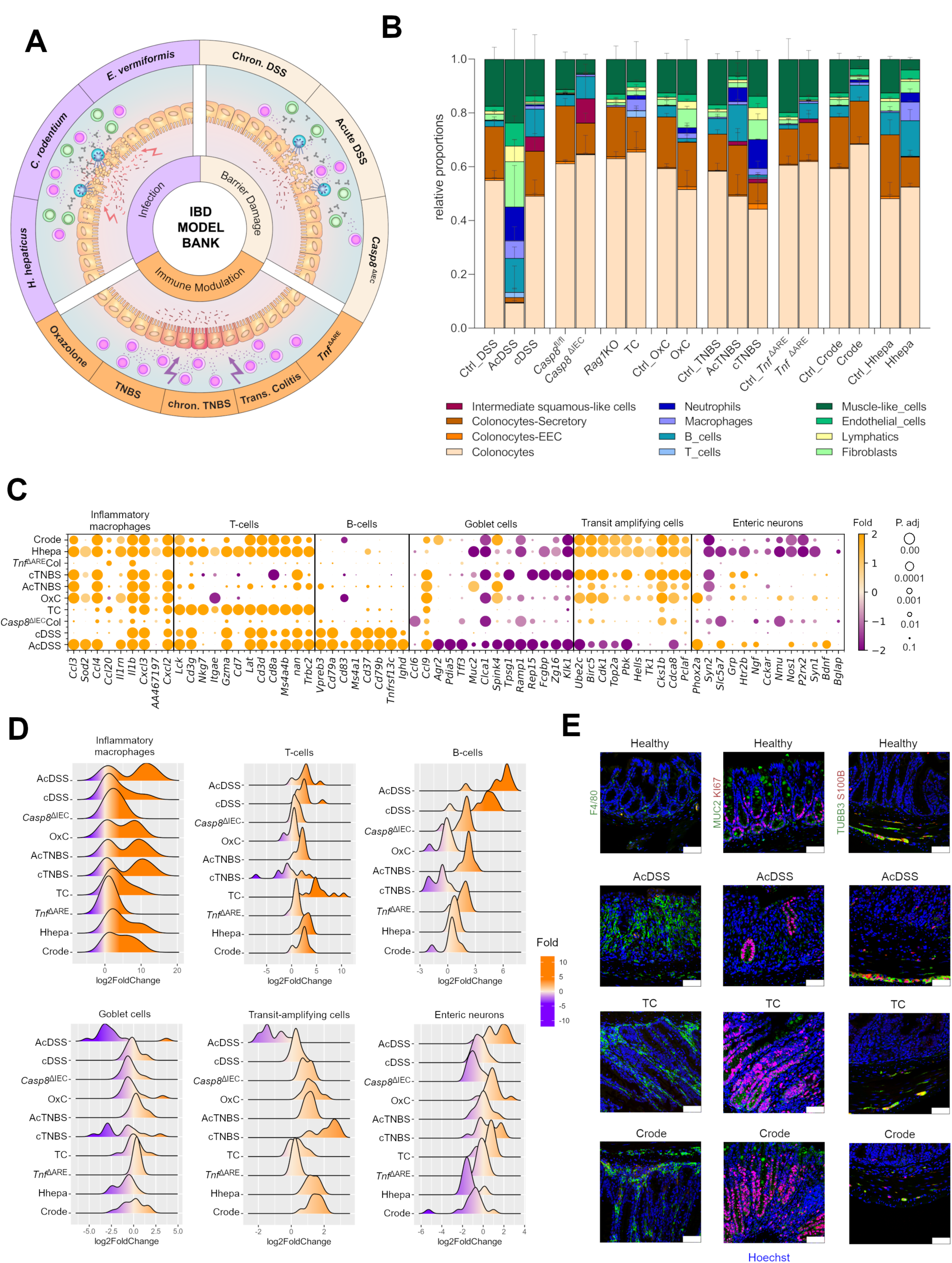
Changes in tissue cell composition across mouse models of IBD. **A**) Scheme depicting the mouse model data bank for preclinical gut inflammation, highlighting the distribution of the broad categories: infection, barrier damage, and immune modulation **B**) Deconvolved cell type signatures across colonic models. Inflamed samples are shown next to controls. The columns Ctrl_OxC and Ctrl_Crode are shared reference controls for the two respective models. **C**) Bubble plot showing fold change and adjusted p values of selected cell type markers across the colonic mouse models. **D**) Ridge plots of the fold change of selected cell type marker genes across the colonic mouse models. **E**) Immunofluorescence staining for markers for macrophages (green, first column), goblet cells (Green, second column), transit amplifying cells (red, second column), enteric neurons (green, third column), and enteric glia (red, third column) across the indicated models from each category.

Next, we created a publicly accessible transcriptomic database and webtool denoted SEPIA (http://trr241.hosting.rrze.uni-erlangen.de/SEPIA/), to explore various aspects of each model, enabling user friendly comparative analyses (Supplemental Fig. S1C). The anatomical location, rather than the inflammatory trigger, accounted for most variation, prompting separate analyses of colonic versus ileal models (Supplemental Fig. S1D). We used scRNA-Seq datasets [14] and deconvolution algorithms [15] to generate signature matrices and extract cellular compositional features from bulk transcriptomes. In inflamed mice, higher immune cell proportions were evident, with distinct cellular compositions across models, including macrophage and neutrophil infiltration, colonocyte subtype loss, and stromal population impacts concurrent to histological and endoscopic findings of epithelial erosion, hypertrophy, and enhanced granularity (Fig. 1B, Supplementary Fig. S1A-B, and Table ST2).

New and unexpected insights included the detection of squamous-like epithelia in cDSS and *Casp8*^ΔIEC^Col, muscle-like cell reduction in AcTNBS, Hhepa, and Crode, and elevated B-cell signatures in AcDSS, cDSS, and AcTNBS models (Fig. 1B, Supplemental Table ST2). A poor representation of certain cell types such as enteric neuroglia, prompted us to take a curated approach for validating the deconvolution results. For this, we analysed marker gene expression representing specific cell populations (Fig. 1C-D, Supplemental Fig. S2A-B). These analyses confirmed the deconvolution data such as loss of secretory goblet cells in AcDSS, AcTNBS, Hhepa and Crode models, and revealed new findings such as an elevation in the transit amplifying (TA) cell markers in the Hhepa, Crode, and cTNBS colitis models, correlating with epithelial hypertrophy (Fig. 1C-D, Supplemental Fig. S1B). Furthermore, we detected that enteric neuronal marker levels contrasted strongly between infectious versus the AcDSS and OxC models (Fig. 1C-D). An interesting inverse correlation was detected between B-cell versus TA-cell markers, and a similar inverse trend emerged between enteric neuron versus TA-cell markers (Fig. 1D). Corresponding trends at the protein level confirmed our transcriptomic findings (Fig. 1E).

In small intestinal models, we observed model-specific immune cell marker changes (Supplemental Fig. S2A, B). Strikingly, the *Tnf*^ΔARE^Ile model exhibited a strong inverse trend between T-and B-cell markers and was the only model with strong B-cell marker expression. Similar to colonic models, a negative correlation between the B-cell and TA-cell markers was also observed. Goblet cell markers were repressed in all small intestinal models, matching histological findings (Supplemental Fig. S1A, S2A, B). Interestingly, the *Casp8*^ΔIEC^Ile model, showed only mild macrophage elevation, T-cell suppression, and no B-cell alteration despite histological inflammation (Supplemental Fig. S1A, S2A, B).

Our analyses demonstrate that transcriptomic tissue alterations largely align with histological and preclinical metrics, revealing significant differences in cellular architecture within and between model categories.

### Common and unique transcriptomic signatures from diverse models simulate IBD pathways

After performing correlative analyses to directly compare transcriptomes between and within each category (Supplemental Fig. S3A, B and Supplementary information), we evaluated gene ontology (GO) similarities among commonly co-expressed genes in colonic (Fig. 2A, upregulated; 2E, downregulated) and small intestinal models (Fig. 2B, upregulated; 2F, downregulated). We performed ontology-based semantic similarity analysis to determine functional attributes between these genes [16, 17]. Commonly upregulated genes in colonic models confirmed expected inflammatory responses, mainly including: innate immune response, chemotaxis, and response to bacteria (Fig. 2A, C). In contrast, small intestinal models showed upregulation in rRNA metabolic process, RNA processing, and telomere organization (Fig. 2B, D).

**Figure 2.**
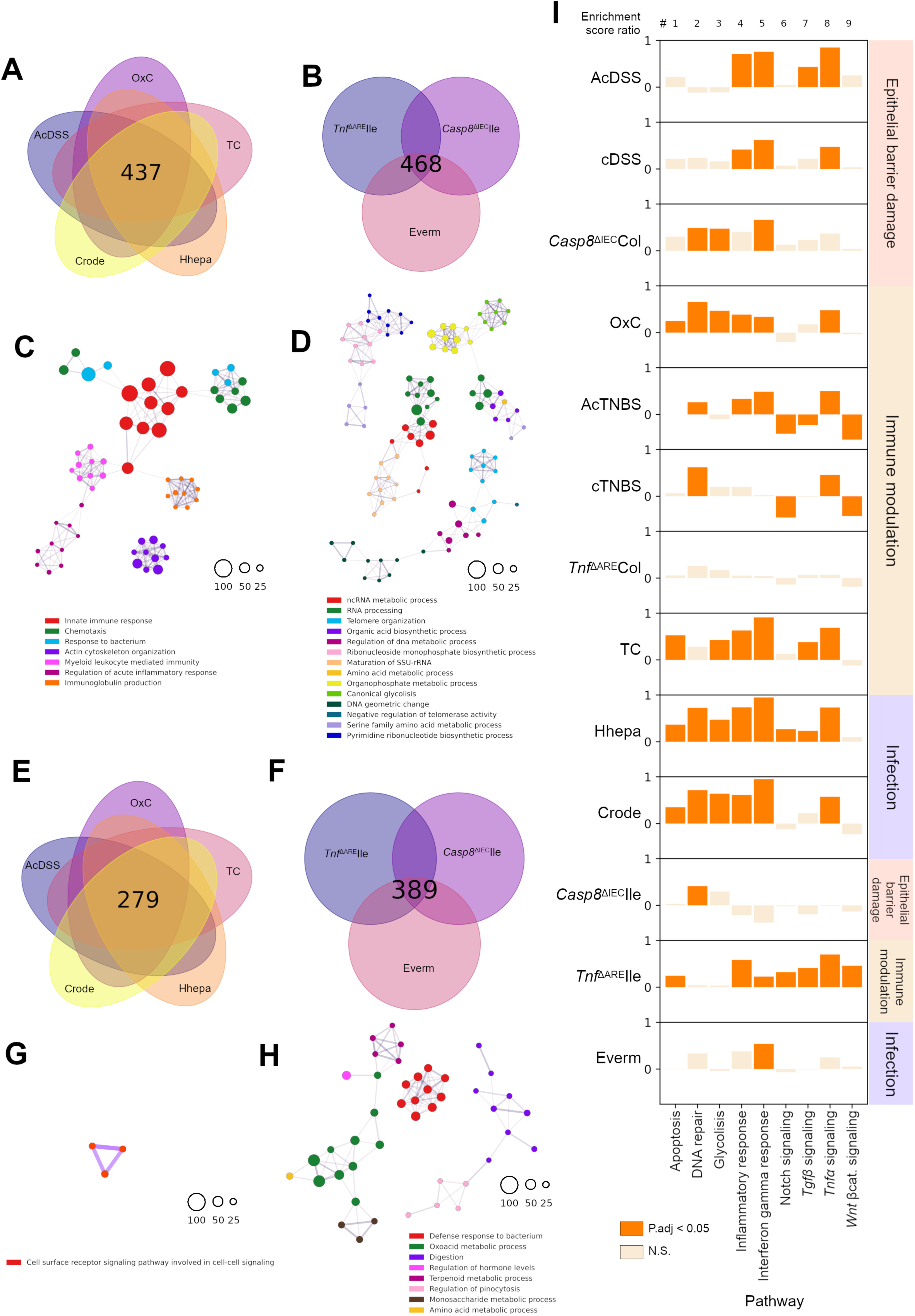
Regulatory commonalities and differences between mouse models. **A**) Venn diagram showing the common upregulated genes between the indicated colitis models and **B)** those between ileitis models. **C-D**) Semantic similarity networks from the enriched GO terms observed in the upregulated gene set from the common datasets defined in A and B respectively. **E**) Venn diagrams showing the common downregulated genes between the indicated colitis models and **F)** those between the ileitis models. **G-H**) Term similarity network of the enriched GO terms observed in the downregulated gene set from the common datasets defined in E and F respectively**. I**) Variation in the relative expression of deconvolved hallmark pathways per mouse model showing variation profiles across the model dataset.

The 279 commonly downregulated genes in colonic models (Fig. 2E) didn’t enrich against known ontology terms, except for Wnt signalling (Fig. 2G). In stark contrast, the 389 commonly downregulated genes in the ileal models (Fig. 2F), were enriched for alpha-defensins indicating suppressed antimicrobial responses (Fig. 2H). Detailed ontology terms, adjusted p-values and kappa coefficients are provided in Supplemental Table ST3. In addition to the shared genes and processes, we also analysed the unique genes specific to each model identifying model-specific ontologies (Supplemental Fig. S4A-D; Supplemental Table ST3).

Next, we investigated the relationship between models and hallmark pathways associated with IBD using gene set variation analysis (GSVA) coupled with limma-based estimation of differences [18, 19, 20]. Nine distinct MySigDB hallmark pathways relevant to inflammation and epithelial homeostasis were selected [21, 22]. The enrichment scores for specific pathways were highly similar for models within each category. In colonic barrier damage models, pathways like inflammatory response, IFNγ response, and TNF signalling were robustly activated. Whereas in the colonic immune modulation category, DNA repair and TNF signalling pathways were activated (Fig. 2I). Crode and Hhepa infection models showed concordance in upregulated pathways, except for Wnt-β-catenin signalling, which was reduced in Crode but unaffected in Hhepa. Notch and TGBβ signalling were upregulated in Hhepa but not in Crode (Fig. 2I). The AcTNBS and OxC models showed reduced Notch and Wnt-β-catenin signalling (Fig. 2I). It was striking that the TC model of immune modulation showed pathway enrichment similar to colonic infection models, with significant enrichment in most pathways, including TNF signalling (Fig. 2I). Among the small intestinal models, the TNF signalling reached significance only in the *Tnf*^ΔARE^Ile model, with a trend in the Everm, but not the *Casp8*^ΔIEC^Ile model (Fig. 2I). The IFNγ response pathway was significantly enriched among all three small intestinal models (Fig. 2I). These analyses provide a reference for selecting appropriate models for investigating specific pathways in IBD.

### Gene co-expression modules in mouse models of intestinal inflammation

To circumvent the limitations arising from pathway annotation and database selection, we employed weighted correlation network analysis (WGCNA) [23] to identify gene co-expression modules across our mouse model datasets. The 16 colonic and 11 ileal modules of co-expression, thus identified (Fig. 3A, B; Supplemental Table ST4-5) demonstrated the expected co-expression of specific modules in both colonic as well as ileal models. For instance, modules ME11 and ME4, respectively, were identified in both locations and consisted of GO biological processes related to “inflammatory responses” (Fig. 3C, D). Similarly, the modules ME3 (colon) and ME1 (ileum) exhibited predominantly negative Δ eigenvalues and were composed of ontologies related to “carboxylic acid processes” (Fig. 3C, D; Supplementary table ST6).

**Figure 3.**
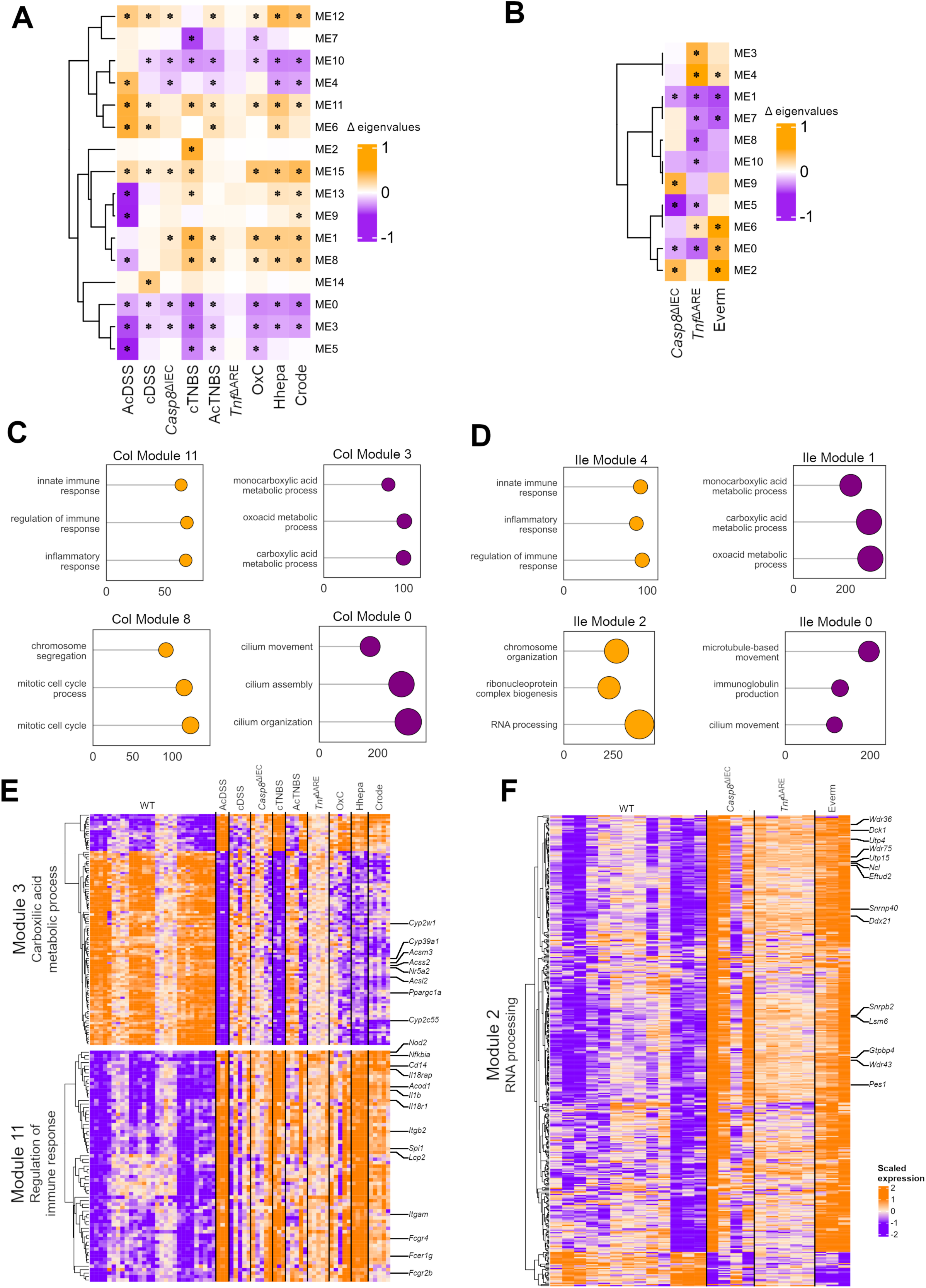
Weighted Gene Co-expression Network Analysis (WGCNA) across the mouse model dataset. **A-B**) Variation observed in the modules obtained from the WGCNA applied to colitis and ileitis samples from the mouse model dataset. Changes labelled with IZ indicate post-limma p values of < 0.05. **C-D**) Top three Gene Ontologies enriched in selected modules from the WGCNA of colitis (C) and ileitis (D) respectively. **E-F**) Heatmap of the enriched genes from the selected WGCNA modules from the colitis and ileitis mouse models, respectively.

Interestingly, the 437 colonic (Fig. 3A) and 468 ileal (Fig. 3B) commonly upregulated genes identified earlier, fell into the ME11 (colon) and ME4 (ileum) modules, respectively. These modules consisted of immune regulatory ontologies with genes such as *Nod2*, *Nfkbia, Il1b*, *Il18r1*, *Lcp2*, *Itgb2*, among others (Fig. 3C-E; Supplemental Table ST4-6).

Moreover, our analysis revealed that ME3 (colon) and ME1 (ileum) consisted of ontologies belonging to the carboxylic acid metabolic process, with notable regulatory genes associated with immune suppression. For example, these modules included the gene *Nr5a2*, which encodes for LRH-1, a protein involved in the generation of immunomodulatory corticosteroids. They also included the genes *Cyp2w1*, *Cyp39a1*, and *Cyp2c55,* which control cholesterol catabolism and supply for the synthesis of corticosteroids [24]. It is well established that both host-derived and commensal carboxylic acid species are known to regulate immune responses. Interestingly, the fact that the same modules also contain genes that regulate the supply of monocarboxylic acids, including, for example, *Acsm3*, *Acss2, Ppargc1a*, and *Acsl2,* which are involved in the control of metabolic immunomodulation (Fig. 3C, E; Supplemental Table ST4,6) [25, 26, 27]. It was interesting that the same genes representing these pathways were also found to be repressed in human IBD, as described later in the following sections (Fig. 4C, D). This suggests that a generalized suppression of a co-regulated network of endogenous immunosuppression across inflamed tissues occurs, regardless of the initial trigger of inflammation or its location.

**Figure 4.**
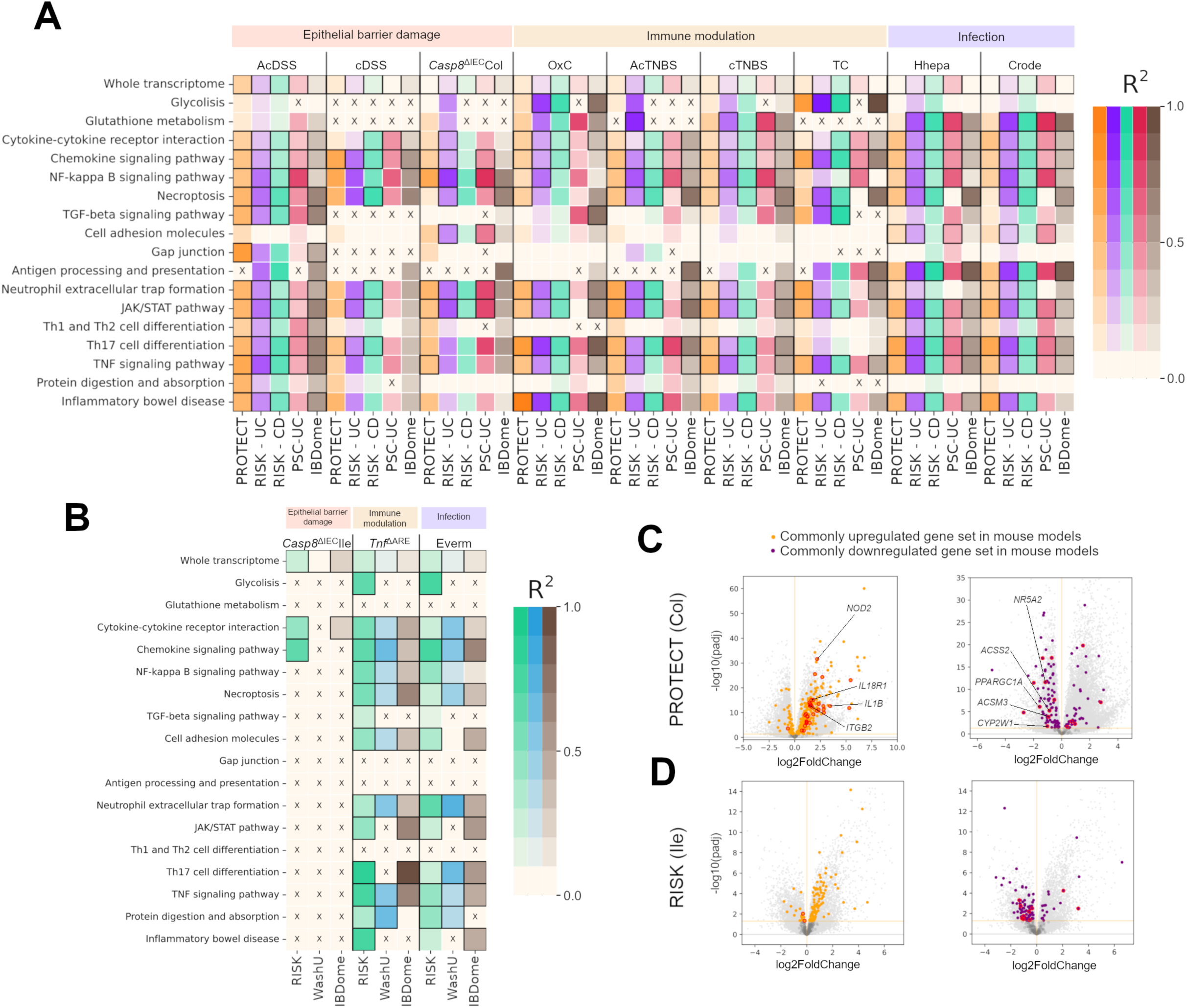
Translation of mouse model features to human IBD. **A-B**) Correlation sets for selected ontologies between (A) multiple mouse colitis models versus human UC cohort transcriptomes and (B) mouse ileitis models versus human CD cohort transcriptomes. Colour intensity represents the R^2^ value. Framed cells denote a significant p-value in the regression test (p < 0.05). Cells marked with an [X] had less than 10 genes in common and the regression was not performed. **C**) Volcano plots of the indicated UC patient cohort where (left) orthologs from the common upregulated genes from colitis mouse models and (right) downregulated genes are shown. All gene commonly regulated among mouse colitis models are highlighted and those belonging to the WGCNA mouse colitis modules ME11 and ME3 are labelled in red **D**) Volcano plots of the indicated CD patient cohort where (left) orthologs from the common upregulated gene set from ileitis mouse models and (right) downregulated genes are shown. All genes commonly regulated among mouse models are highlighted and those belonging to the WGCNA mouse ileitis modules ME2 and ME1 are labelled in red.

In addition to the aforementioned common modules, tissue-and model-specific co-regulatory gene ontologies with opposing Δ eigenvalues were also identified. For example, ontologies pertaining to “neuronal processes” were observed in ME4 (colon), which exhibited a specific and inverse correlation in the AcDSS model (Fig. 3A). Similarly, in the ileal models, ME2 consisted of unique gene ontologies for “RNA processing” with positive Δ eigenvalues for the genes *Wdr36*, *Dck1*, *Utp4*, and *Wdr75*, which control ribosomal RNA processing. These genes play a role in the context of p53 stress-responses [28, 29, 30, 31] (Fig. 3F; Supplemental Table ST5-6).

### Conserved and divergent co-expression ontologies between preclinical models and IBD patients

Comparative transcriptomics enables contrasting patient and preclinical models, facilitating the identification of conserved disease mechanisms and preclinical testing [32]. Using public and our own IBD-patient transcriptomes (Supplemental table ST7), we compared them with our mouse model data, analysing commonly regulated IBD relevant ontologies [33, 34]. This revealed a number of interesting correlations for specific IBD-relevant KEGG pathways, stratified by disease location: colonic (Fig. 4A) and ileal (Fig. 4B). The core pathways driving inflammation, included chemokine signalling, cytokine receptor signalling, the JAK/STAT pathway, and TNF signalling, with higher R^2^ values across model-human cohort pairs (Fig. 4 A, B, and Supplemental Table ST8).

Certain KEGG pathways showed interesting correlations between model–cohort pairs. For instance, in specific IBD cohorts, glutathione metabolism, cell adhesion, and antigen processing and presentation exhibited a high R^2^ and met significance thresholds against infectious colitis models (Fig. 4A). However, among the small intestinal models, several orthologs did not reach the expected expression-enrichment thresholds (Fig. 4B, ‘x’ marks). A distinctive correlation pattern was observed for the pathway of gap junctions in the AcDSS model versus the PROTECT and IBDome patient cohorts. The Th1-Th2 differentiation pathway showed significant R^2^ for the Hhepa and AcDSS models versus most IBD cohorts (Fig. 4A, Supplemental table ST8).

The Th17 differentiation pathway is an important player in the pathogenesis of IBD. This inflammatory pathway of Th17 cell differentiation showed an interesting trend, with significant to high R^2^ values. Among the colonic models, this pathway was significantly correlated in most of the model-cohort pairs, except the *Casp8*^ΔIEC^Col and TC models (Fig. 4A). Similarly, in the ileal models, Th17 cell differentiation was significant in most model-cohort pairs (5/6) where selection thresholds were met (Fig. 4B).

The IBD cohorts in our study are broadly divided into paediatric (PROTECT and RISK) and adult (PSC-UC, WASH-U, and IBDome). We identified specific correlations unique to paediatric cohorts, such as glycolysis in the *Tnf*^ΔARE^Ile and Everm models; TGFβ signalling in the TC and *Tnf*^ΔARE^Ile models; JAK/STAT signalling in the TC model; and Th17 cell differentiation in the cDSS model (Fig. 4A, B).

Interestingly, several of the genes from colonic modules ME11 and ME3 were also concordantly regulated in the PROTECT (UC) patient transcriptomes with similar magnitudes and directions (Fig. 4C). Most genes commonly regulated among ileal models (Fig. 2B, F) were also regulated in the RISK (CD) patient cohort (Fig. 4D). This regulatory overlap between the mouse models and CD patients exhibited a conserved direction and magnitude. Furthermore, these commonly regulated genes from the mouse ileal models (Fig. 2B, F) were represented on the ileal co-expression modules, ME1 and ME2 (Fig. 3D, F). However, the specific ontology associated with RNA processing was not affected in the human RISK cohort (Fig. 4D, lack of highlighted datapoints). Thus, comparative transcriptomics between IBD patient samples and preclinical mouse models revealed conserved disease mechanisms that replicate in at least one of the mouse models in a given category, including key pathways like chemokine signalling, JAK/STAT, and Th17 cell differentiation. On the other hand, regulation of specific pathways correlates better between a given mode-cohort pairs. For example, glycolysis across the immune modulation and the infectious categories and glutathione metabolism and antigen presentation across the infectious categories. These findings highlight shared and unique regulation of key IBD relevant pathways in model-cohort pairs.

### Specific conserved ontologies have unique patterns of expression in IBD-associated cell clusters

Recent single-cell RNA sequencing (scRNA-Seq) data has revealed new cellular states and subsets emerging during gut inflammation and exhibiting non-canonical expression profiles. For example, Li *et al*. identified a new specialized IEC subtype with the capacity to regulate mucosal immunity and drive human CD [35]. To ascertain whether specific inflammation-associated co-expression modules that we identified, can be observed in non-canonical cell subsets emerging in human IBD, we analysed our data against scRNA-Seq studies from IBD patients.

Indeed, analysis of UC patient-scRNA-Seq datasets [36] revealed the emergence of cross-functional states of TA, stem, and immature enterocytes highly expressing genes in the colonic module ME11 enriched for the GO “regulation of imflammatory process”. This GO is canonically ascribed to immune cell lineages (Fig. 5A). In addition, M-cells, which are critical for host defence, also showed an upregulation of this GO class in inflamed UC tissues (Fig. 5A).

**Figure 5.**
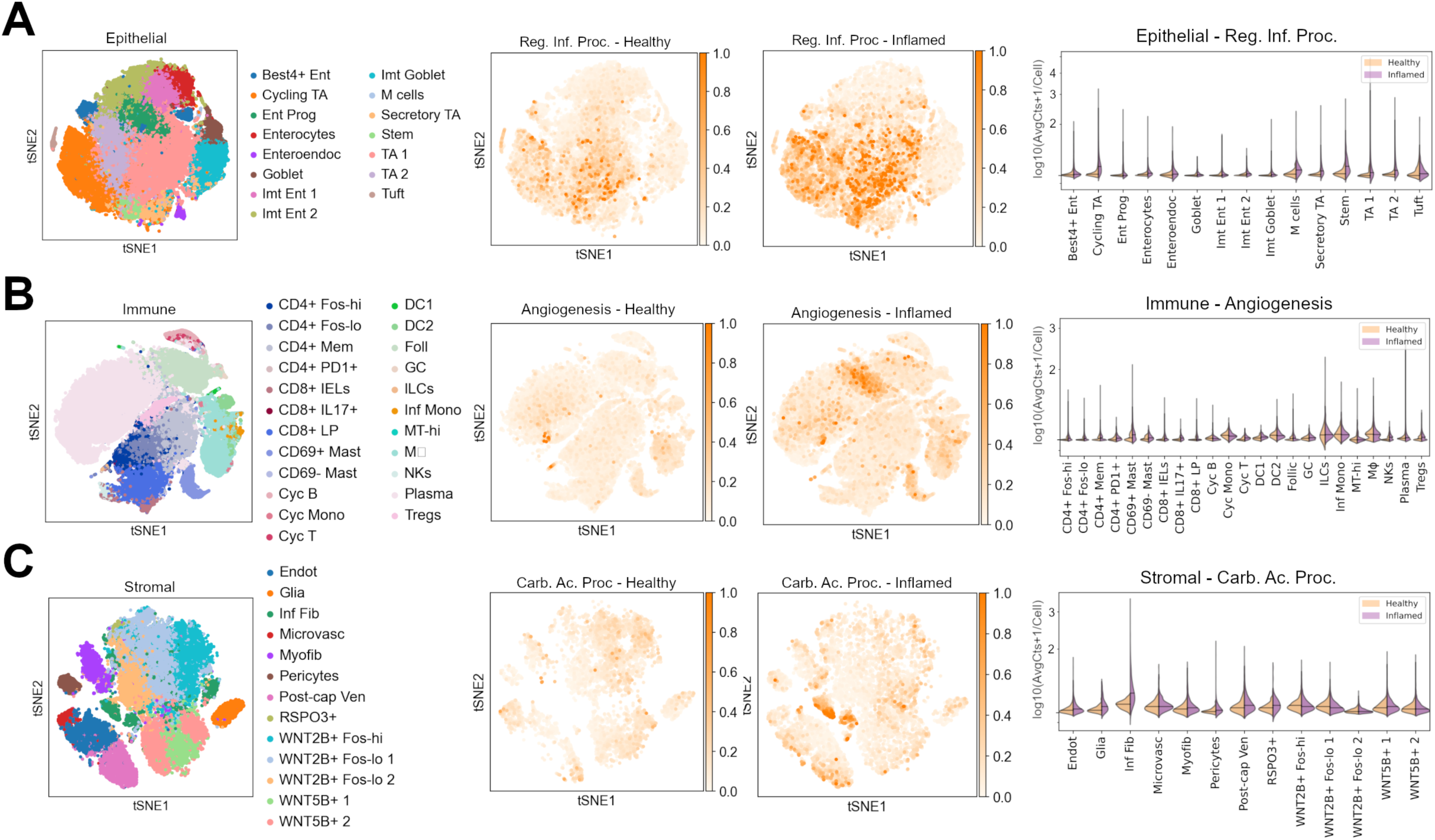
Conserved ontologies are expressed by unique IBD-associated single cell clusters. scRNA-Seq clusters of **A)** epithelial, **B)** immune, and **C)** stromal compartments from Smillie *et al.* [36]. Left: t-SNE clusters, middle: t-SNEs segregated by patient health where colour represents the average gene expression of selected pathways from the WGCNA modules, and right: violin plots showing the changes of the average expression in each of the annotated clusters.

Another novel cellular state, specific to inflamed patients was a plasma cell subcluster. This subcluster displayed an elevated expression of genes enriched for the GO term “angiogenesis”, as identified in the colonic module ME5 (Fig. 5B). Additionally, an elevated level of expression of the GO term “angiogenesis” was observed on the CD69^+^ mast cells cluster specifically in inflamed UC tissues (Fig. 5B). Surprisingly, a recently discovered novel stromal cluster of inflammatory fibroblasts and subclusters of endothelial and post-capillary venule endothelial cells exhibited an elevated expression of genes in colonic module ME3 enriched for the GO “carboxylic acid metabolic process” (Fig. 5C). We also performed an extended module-by-cell type analysis at the gene level for each colonic cell lineage (epithelial, immune, and stromal) and stratified by disease status. We observed a notable degree of cell type specificity in the expression of genes from various modules such as ME5 (goblet), ME8 (cycling TA), and ME14 (M cells), in addition to other differences in the average expression of modules based on the disease status. For example, expression of genes from ME14 with top enrichments related to squamous-like cells was highly enriched in inflamed UC M-cells (Supplemental Fig. S5A; Supplementary Table ST6, 9).

Among the ileitis mouse models, we identified “RNA processing” as an enriched GO class from the module ME2 (Fig. 3F), which did not align on any of the IBD cell clusters (Supplemental Fig. S6A). Moreover, the magnitude of expression for this GO class was comparable across clusters from the scRNA-Seq data of CD ileal samples [37] (Supplemental Fig. S6A, B). However, we detected other modules derived from ileitis mouse models that exhibited cellular-level alterations when compared between ileal CD scRNA-Seq clusters (Supplemental Fig. S6B; Supplemental Table ST9).

Collectively our analyses highlight how conserved expression of specific ontologies results from the emergence of specific cellular states during inflammation in the gut. Since the modules analysed here were drawn from the commonly regulated transcripts across mouse models, it is apparent that different modalities of inflammation triggering evoke similar cell state changes in IBD.

### Inflammatory impacts on cellular substructures are conserved in specific model-cohort pairs

Aberrant transcript expression and physical organization of IEC substructures, such as microvilli, has been reported to affect patients with CD suggestive of epithelial dysfunction [38]. Interestingly, analysis of our preclinical model data had also identified a common co-expression module, ME0, containing several ontologies related to “cilium” (Fig. 3C, D). Indeed, WGCNA modules with negative Δ eigengene scores for ontologies related to “cilium” were enriched in adult colonic PSC-UC (ME1) cohort, almost reaching significance [39] (Fig. 6A-C). A module ME0, consisting of ontologies related to “cilium” was also detected in the RISK ileum CD cohort, but it did not show any significant enrichment (Fig. 6A-C). We recapitulated the original findings from the WashU ileum CD cohort where ciliary dysfunction was first reported, validating the analysis approach (Fig. 6A-C). Confirming this, scanning electron micrographs of the epithelia from patients in our study cohort revealed microvilli damage and their disrupted organization, supporting the finding that this feature is not only restricted to CD, but can also contribute to epithelial dysfunction in UC (Fig. 6D-E). However, it is noteworthy that none of the paediatric IBD cohorts, including RISK and PROTECT, had modules containing enrichments for “cilium” or related ontologies indicative of an age dependency in this subcellular dysfunction. Moreover, cilium and related ontologies were significantly depleted in specific mouse models including the colitis models Crode, cTNBS, and AcTNBS and the ileitis models Everm and *Tnf*^ΔARE^ (Fig. 6F-G). Our data indicate that damage to epithelial microvilli is common in CD as well as UC and affects adult more than paediatric IBD patients and this feature can be modelled in specific preclinical mouse models of intestinal inflammation.

**Figure 6.**
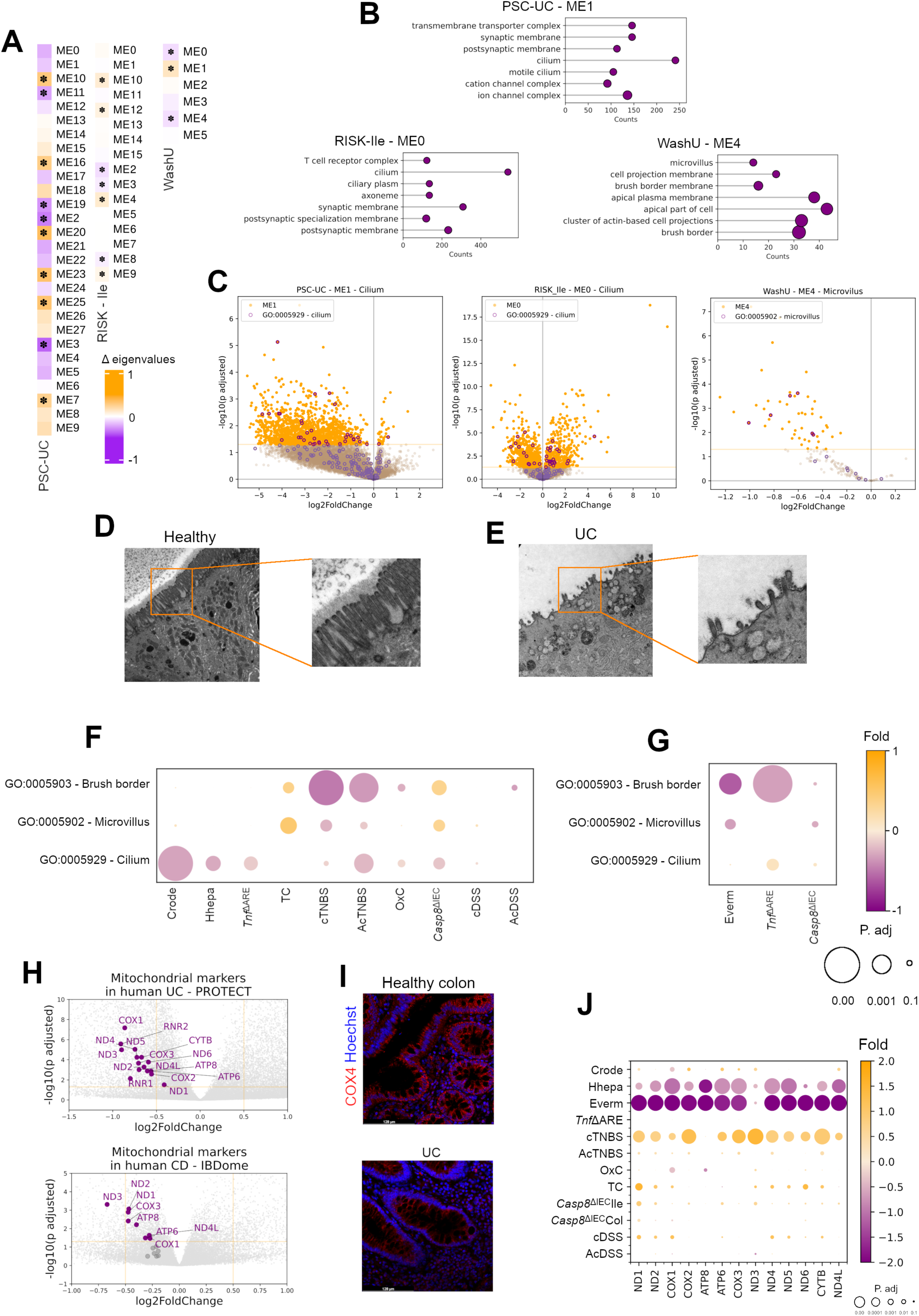
Microvilli dysfunction and mitochondriopathy observed in IBD translate to selected mouse colitis and ileitis models. **A)** Variation observed in the mouse colitis and ileitis WGCNA modules applied to the indicated human cohorts. Significant changes labelled with * indicating p values of < 0.05 in the differential analyses. **B)** Top 7 ontologies enriched in selected modules from the analysis in A). **C)** Volcano plots of the indicated human cohorts with genes from the selected WGCNA modules highlighted in orange and those from the specific enriched ontologies from B) in purple. **D)** Electron micrographs of the intestinal epithelial cells from healthy individuals. **E)** Electron micrographs of epithelial cells from UC patients **F-G)** Bubble plot showing the variation in the relative expression of selected ontologies from the mouse colitis F) and ileitis G) models. **H)** Volcano plots from the indicated UC and CD patient transcriptomes with mitochondrially expressed genes. **I)** Immunofluorescence staining for mitochondrial marker COX4 in healthy individuals and UC patient colonic samples. **J)** Bubble plot showing the log2 fold change in the expression of mitochondrially encoded genes across all mouse models of colitis and ileitis.

Another subcellular organelle impacted in IBD are mitochondria. A recent study demonstrated how mitochondriopathy contributes to disease severity and treatment response in the UC PROTECT cohort [40, 41]. We replicated these findings in our in-house cohort where the expression levels of multiple mitochondrially encoded genes and the protein levels of COX4 were reduced in the UC patient samples (Fig. 6H-I). Interestingly, extrapolation of these findings to our mouse model data, showed that the mitochondrial changes were most closely modelled in the infectious category, Everm and Hhepa (Fig. 6J). It is noteworthy that in contrast to the infectious mouse models, the cTNBS colitis model showed an inverse correlation with respect to the expression of mitochondrially encoded genes versus that in UC patients.

Overall, our data highlight the conserved and divergent regulation of genes and pathways in preclinical intestinal inflammation models and clinical IBD cohorts. Utilizing these as a tool for pre-screening while targeting IBD processes will enable guided discovery efforts.

## Discussion

Preclinical modelling of diseases to understand the underlying mechanisms of chronic inflammation are hindered by the poor translatability of results. A common challenge is selecting an appropriate model to study the pathway and mechanism in question. Given the diversity of models and different modes of inflammation induction, this choice becomes even more complex. Our data summarize how comparative transcriptomic analyses can elucidate shared and model-specific regulatory features and identify translatable components between preclinical models and clinical disease. The core regulatory overlaps, between the disparate preclinical models are well-conserved in clinical IBD. Conversely, each mouse model also has its unique regulatory features which enables the preselection of appropriate models for investigating specific molecular pathways.

In conducting these analyses, we also compared the alterations occurring at the tissue level, tracking cellular changes using deconvolution approaches. Deconvolution is more robust than other methods because it does not rely on preselected cell type markers. These analyses revealed the emergence of specific cell subsets in given mouse models such as the recently reported squamous epithelial cells [14]. The presence of intermediate squamous-like cells was identified in the *Casp8*^ΔIEC^Col, cDSS, and AcTNBS, models. In our data, an interesting inverse association was observed between TA-cells and B-cells, consistent with the recent observation that B-cell expansion impedes epithelial regeneration [42]. Further studies are necessary to elucidate this interplay between epithelial regeneration and B-cells in the context of IBD.

By comparing the analytical findings from the mouse model data with IBD patient datasets, we were able to recapitulate regulatory behaviours of conserved pathways in IBD. We analysed publicly available IBD transcriptomic datasets from the paediatric PROTECT and RISK cohorts [40, 43], alongside biopsy-derived transcriptomes from three patient cohorts: our IBDome UC and CD cohort, a PSC-UC cohort [39], and an adult CD cohort (WashU) [38]. Interestingly, the upregulation of core inflammatory genes was conserved between colonic mouse models and IBD cohorts, while downregulation of immunomodulatory pathways was conserved in the mouse model, PROTECT, and partly in RISK cohorts. Indeed, specific processes were associated with the adult IBD cohorts, which did not recapitulate in the paediatric cohorts. For example, the identification of the downregulation of the GO processes ciliary organization and elevated microvilli damage was conserved among the adult IBD cohorts of PSC-UC and WashU and specific mouse models [38]. Moreover, a mitochondriopathy-like phenotype was observed in the paediatric PROTECT and RISK cohorts that was shared in the infectious colitis models but not in cTNBS exemplifying modelling potential of specific features from given patient cohorts [40, 43].

By superimposing signatures from specific co-expression modules on scRNA-Seq data from immune cell lineages of IBD patients, we identified a cluster of plasma cells with lymphangiogenic functions in the inflamed tissues. These findings are consistent with the recent identification of a new B-cell subset with angiogenic properties by van de Veen *et al*. [44]. The elevated average expression of the carboxylic acid metabolic process in inflammatory fibroblasts was intriguing. The GO class carboxylic acid metabolic process, which controls the supply of a plethora of endogenous metabolites, also implicated in immune cell regulation [45, 46], demonstrated an overall repression in our mouse model data as well as in IBD transcriptomes. Specific stromal clusters from inflamed tissues of UC patients, including inflammatory fibroblasts, post-capillary venule endothelia, and glia, showed an elevation in the expression for genes of carboxylic acid metabolic process. A recent hypothesis suggests that during inflammation and cancer, stromal cells may act as a source of gut-specific immunosuppressive mediators [47, 48, 49]. However, this may also reflect high metabolic demands on these cells, driving the production of large quantities of extracellular matrix molecules or metabolites derived from carboxy acids. Further characterization of this pathway during inflammation and remission in the context of paracrine immunosuppression are warranted.

In summary, we present a resource for the rapid profiling of global transcriptomic alterations in preclinical intestinal inflammation models and provide comparative references against human cohorts. The observations made in our study have important implications for choosing mouse models to investigate disease-driving mechanisms in IBD. Finally, our analyses provide evidence for the composite, overlapping, and conserved program of inflammation in intestinal inflammation, highlight unique model-specific features, and provide a resource for modelling and translation of intestinal inflammation. Further investigation and replication of these findings will help reveal unique molecularly-defined IBD subgroups. This will also enable a guided choice of mouse models, to investigate the intended disease mechanism in light of its regulatory conservation in human disease.

## Materials and methods

Detailed materials and protocols are provided in the supplemental methods section.

## Supporting information

SupplTable1 - Rereferences and information for mouse models

SupplTable2 - 2way ANOVA cellular deconvolution

SupplTable3 - common and exclusive enriched GO ontologies

SupplTable4 - WGCNA module gene composition - colon

SupplTable5 - WGCNA module gene composition - ileum

SupplTable6 - WGCNA enriched GO ontologies

SupplTable7 - Rereferences and information for human cohorts

SupplTable8 - Human-mouse correlation details

SupplTable9 - WGCNA modules in single cell

## Data availability

All of the raw sequencing data generated from this study has been uploaded to the European nucleotide archive via ArrayExpress (https://www.ebi.ac.uk/biostudies/arrayexpress) and is available for download using the following accession numbers (AcDSS, cDSS, and TC: E-MTAB-14306; Everm: E-MTAB-14297; Hhepa: E-MTAB–14316; OxC, Crode: E-MTAB– 14312; *Casp8*^ΔIEC^Col and *Casp8*^ΔIEC^Ile: E-MTAB-14318; AcTNBS, cTNBS: E-MTAB-14329; *Tnf*^ΔARE^Col and *Tnf*^ΔARE^Ile: E-MTAB-14325). We have also created an interactive webtool that enables accessing and browsing of processed data that can be accessed using the following link: http://trr241.hosting.rrze.uni-erlangen.de/SEPIA/ (user login available upon request). Publicly available IBD cohorts can be accessed at the gene expression omnibus https://www.ncbi.nlm.nih.gov/geo/ (accession IDs-PROTECT: GSE109142, RISK CD: GSE57945, and RISK UC: GSE117993) and ArrayExpress (accession IDs-PSC-UC: E-MTAB-7915 and WashU: E-MTAB-5783). Access to the IBDome dataset is available upon request. Publicly available mouse single cell RNA sequencing datasets used in this study can be accessed using the following accession number: GSE168033. Accession for human scRNA-Seq datasets used in this study are UC SCP259 and CD SCP1423.

## Code availability

All code generated in this study has been deposited on the following GitHub repository and is accessible via the following link https://github.com/MiguelGonzalezAcera/SEPIA.

### Acknowledgements

We acknowledge the support provided for animal husbandry and rearing for this work by Eva Wellein and Melanie Ziegler. Parts of this manuscript are under preparation for being submitted as part of the thesis leading to the academic title doctor of philosophy for MGA.

## Funding support

This work was funded by the Deutsche Forschungsgemeinschaft (DFG, German Research Foundation) - TRR241 375876048 (A03, A05, A08, B05, Z03), SFB1181 (C05), and individual grants with project numbers 418055832, 531560612, and 510624836. The project was further supported by CRU 5023 (project number 50474582), CRC 1449-B04 and Z02 (project number 431232613); CRC 1340-B06 (project number 372486779). A.N.H is supported by a Lichtenberg fellowship and “Corona Crisis and Beyond” grant by the Volkswagen Foundation, a BIH Clinician Scientist grant and INST 335/597-1, as well as with the ERC-StG “iMOTIONS” grant (101078069). Moreover, the project was supported by the DFG grant covering sequencing costs to B.S and C.B Projektnummer 418055832. The project was also supported by the Interdisciplinary Centre for Clinical Research (IZKF: A76, A93, J96, and ELAN P120). The authors declare no financial and non-financial conflicts of interest arising from this work.

## Author Contributions

The study was conceived by MGA, JVP, and CB and planned BS, ANH, CB, MN, ZT, and the IBDome Consortium. Data was analysed by MGA, JVP, CP, and GS. The figures were prepared by MB, MGA, and JVP. MGA and JVP wrote the manuscript. Help with manuscript editing was provided by MGA, JVP, ANH, BS, CB, ZT, RGB, KH, LE, MB, and AAK. Data was acquired by CP, GS, LE, RC, RGB, TL, LB, MD, RW, IS, KG, FM, and KK. Data interpretation was performed by MGA, JVP, HL, ANH, and CB. Help with securing funds JVP, CB, MN, BS, ANH, SW, BW, CG, KH, and the IBDome Consortium. Study Supervision JVP, KH, ZT, SW, BW, and CB.

## Other contributing authors

TRR241 IBDome Consortium: Imke Atreya^1^, Raja Atreya^1^, Petra Bacher^2,3^, Christoph Becker^1^, Christian Bojarski^4^, Nathalie Britzen-Laurent^1^, Caroline Bosch-Voskens^1^, Hyun-Dong Chang^5^, Andreas Diefenbach^6^, Claudia Günther^1^, Ahmed N. Hegazy^4^, Kai Hildner^1^, Christoph S. N. Klose^6^, Kristina Koop^1^, Susanne Krug^4^, Anja A. Kühl^4^, Moritz Leppkes^1^, Rocío López-Posadas^1^, Leif S.-H. Ludwig^7^, Clemens Neufert^1^, Markus Neurath^1^, Jay V. Patankar^1^, Magdalena Prüß^3^, Andreas Radbruch^5^, Chiara Romagnani^3^, Francesca Ronchi^6^, Ashley Sanders^4,8^, Alexander Scheffold^2^, Jörg-Dieter Schulzke^4^, Michael Schumann^4^, Sebastian Schürmann^1^, Britta Siegmund^4^, Michael Stürzl^1^, Zlatko Trajanoski^9^, Antigoni Triantafyllopoulou^5,10^, Maximilian Waldner^1^, Carl Weidinger^4^, Stefan Wirtz^1^, Sebastian Zundler^1^

^1^Department of Medicine 1, Friedrich-Alexander University, Erlangen, Germany

^2^Institute of Clinical Molecular Biology, Christian-Albrecht University of Kiel, Kiel, Germany.

^3^Institute of Immunology, Christian-Albrecht University of Kiel and UKSH Schleswig-Holstein, Kiel, Germany.

^4^Charité – Universitätsmedizin Berlin, corporate member of Freie Universität Berlin and Humboldt-Universität zu Berlin, Department of Gastroenterology, Infectious Diseases and Rheumatology, Berlin, Germany

^5^Deutsches Rheuma-Forschungszentrum, ein Institut der Leibniz-Gemeinschaft, Berlin, Germany

^6^Charité – Universitätsmedizin Berlin, corporate member of Freie Universität Berlin and Humboldt-Universität zu Berlin, Institute of Microbiology, Infectious Diseases and Immunology

^7^Berlin Institute für Gesundheitsforschung, Medizinische System Biologie, Charité – Universitätsmedizin Berlin

^8^Max Delbrück Center für Molekulare Medizin, Charité – Universitätsmedizin Berlin

^9^Biocenter, Institute of Bioinformatics, Medical University of Innsbruck, Innsbruck, Austria.

^10^Charité – Universitätsmedizin Berlin, corporate member of Freie Universität Berlin and Humboldt-Universität zu Berlin, Department of Rheumatology and Clinical Immunology, Berlin, Germany

---

**Supplementary figure 1.**
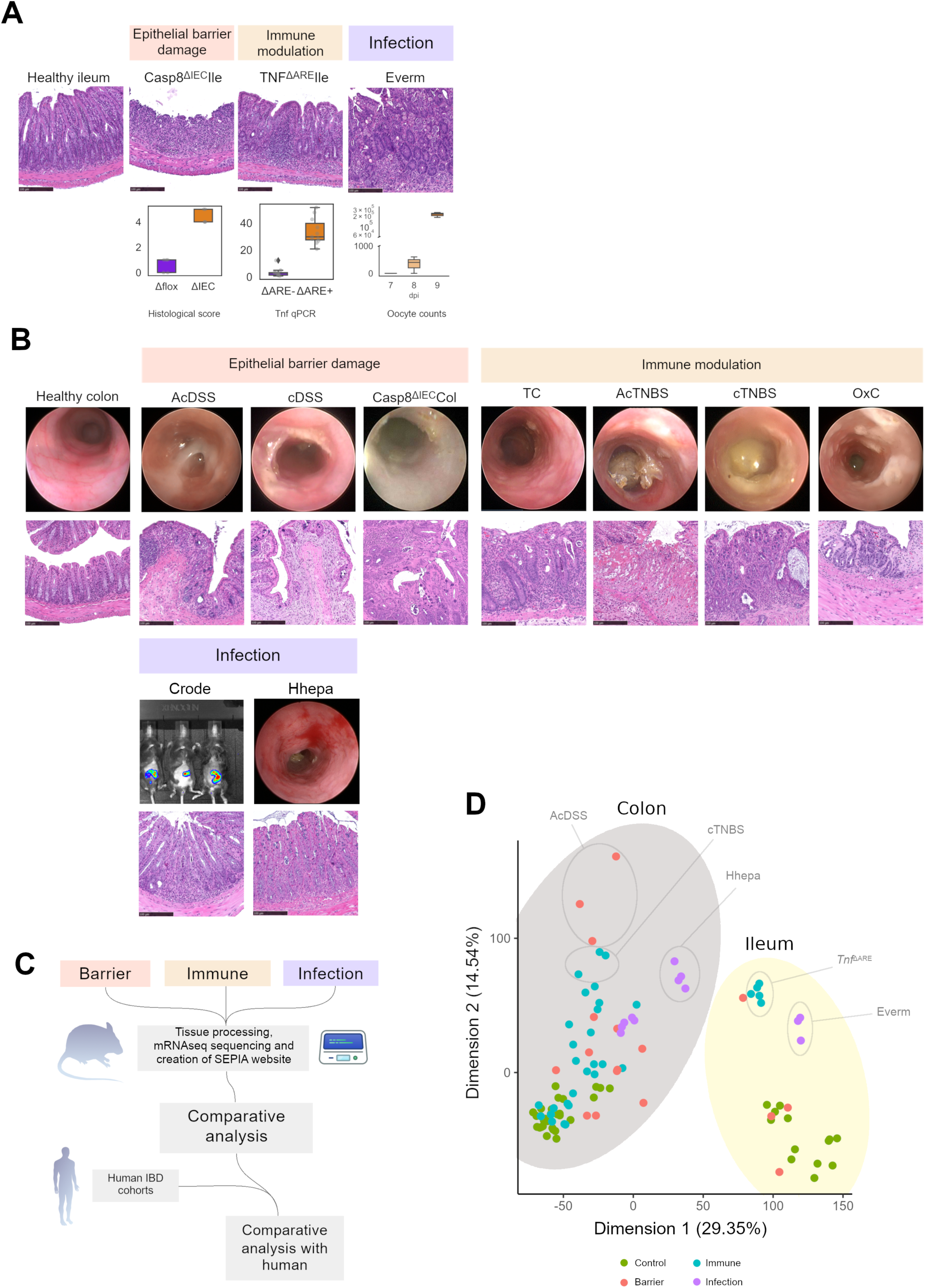

**Supplementary figure 2.**
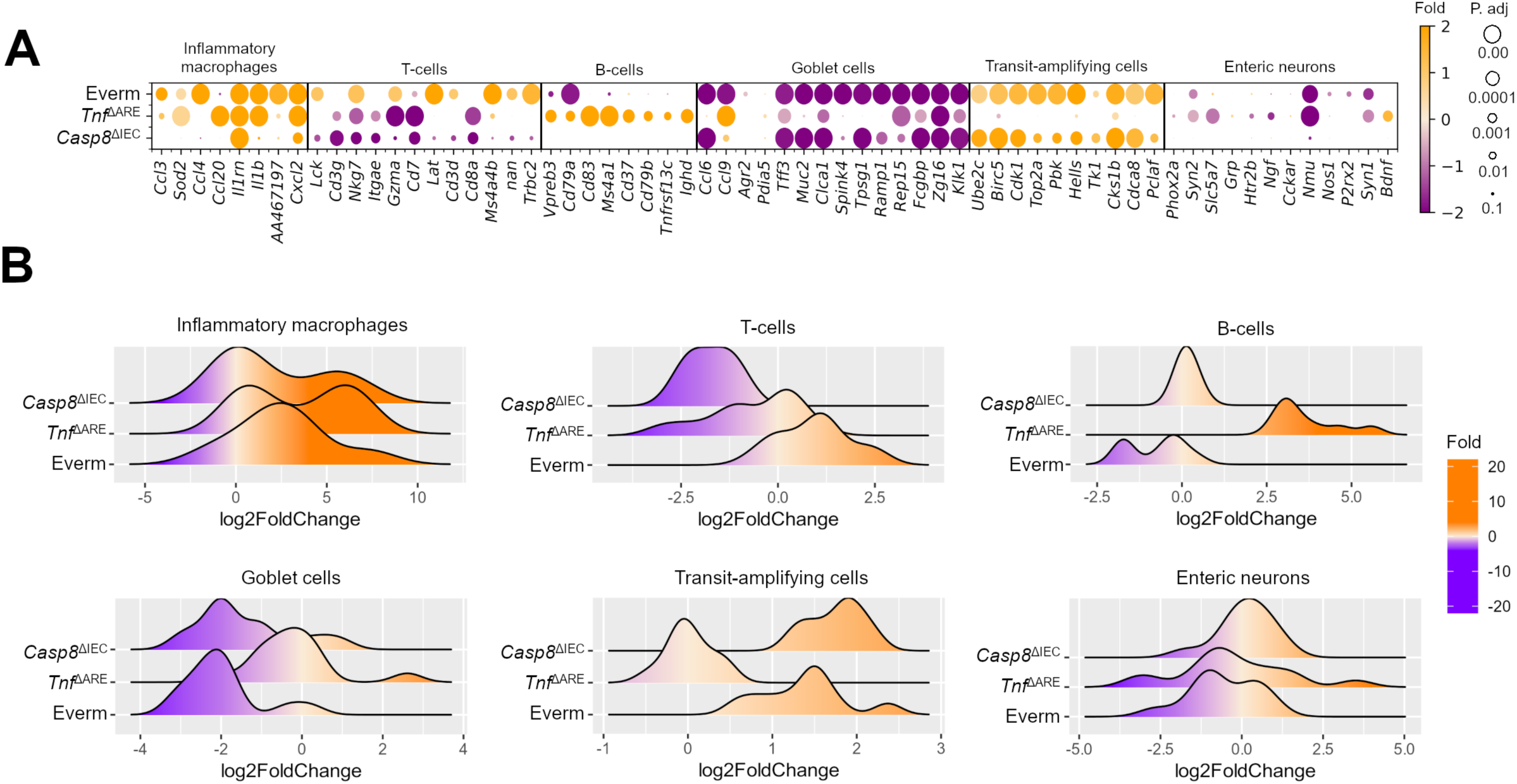

**Supplementary figure 3.**
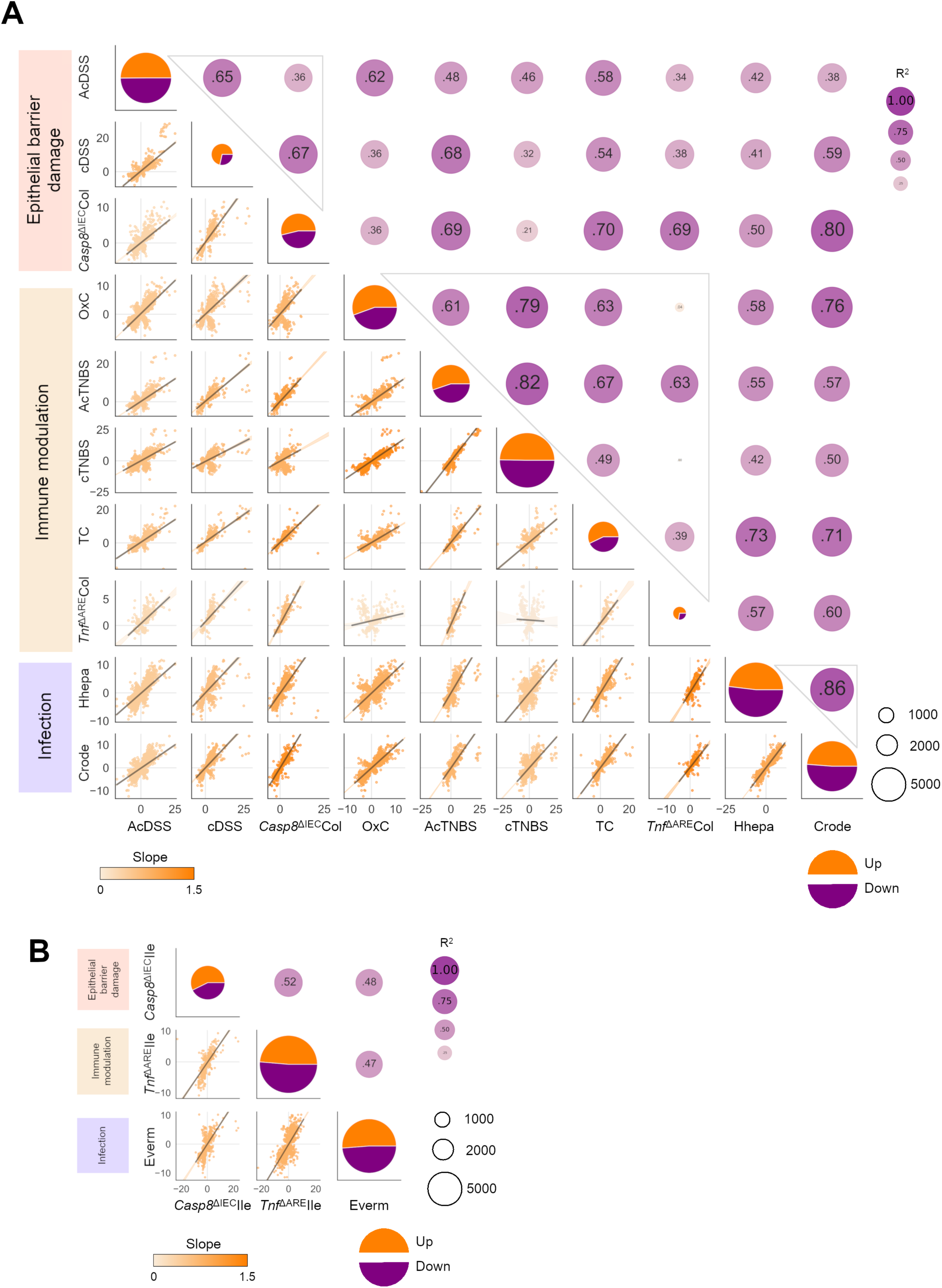

**Supplementary figure 4.**
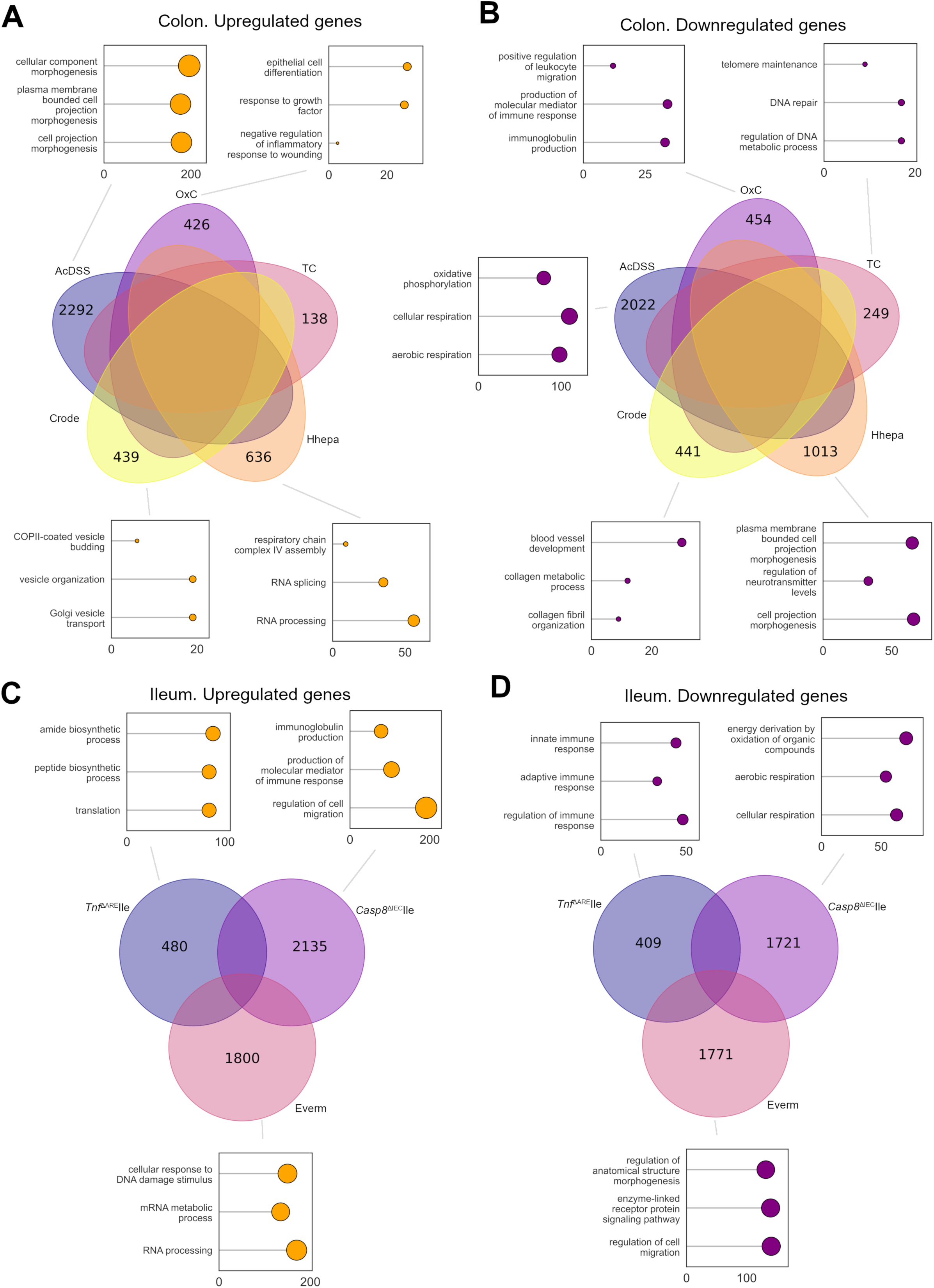

**Supplementary figure 5.**
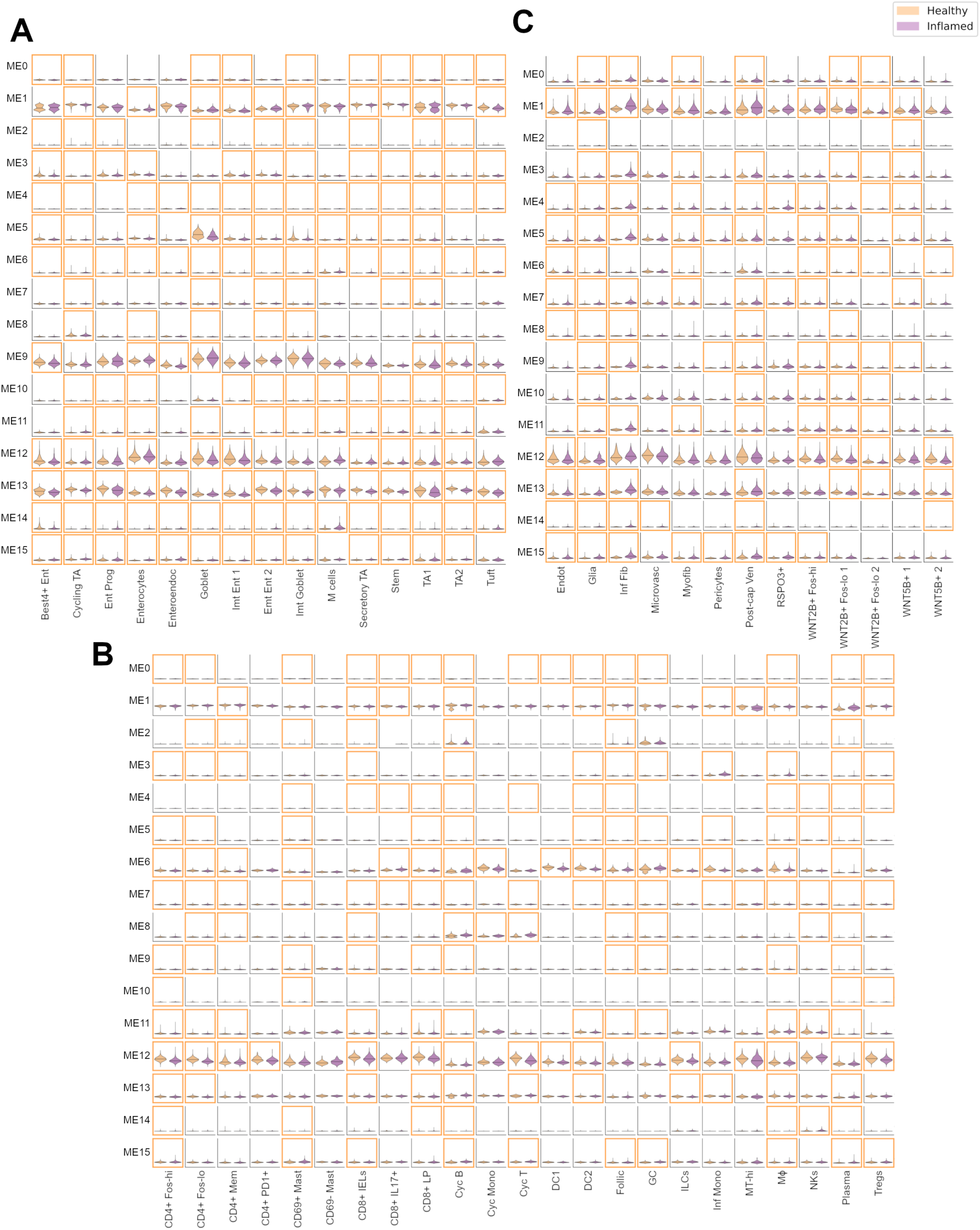

**Supplementary figure 6.**
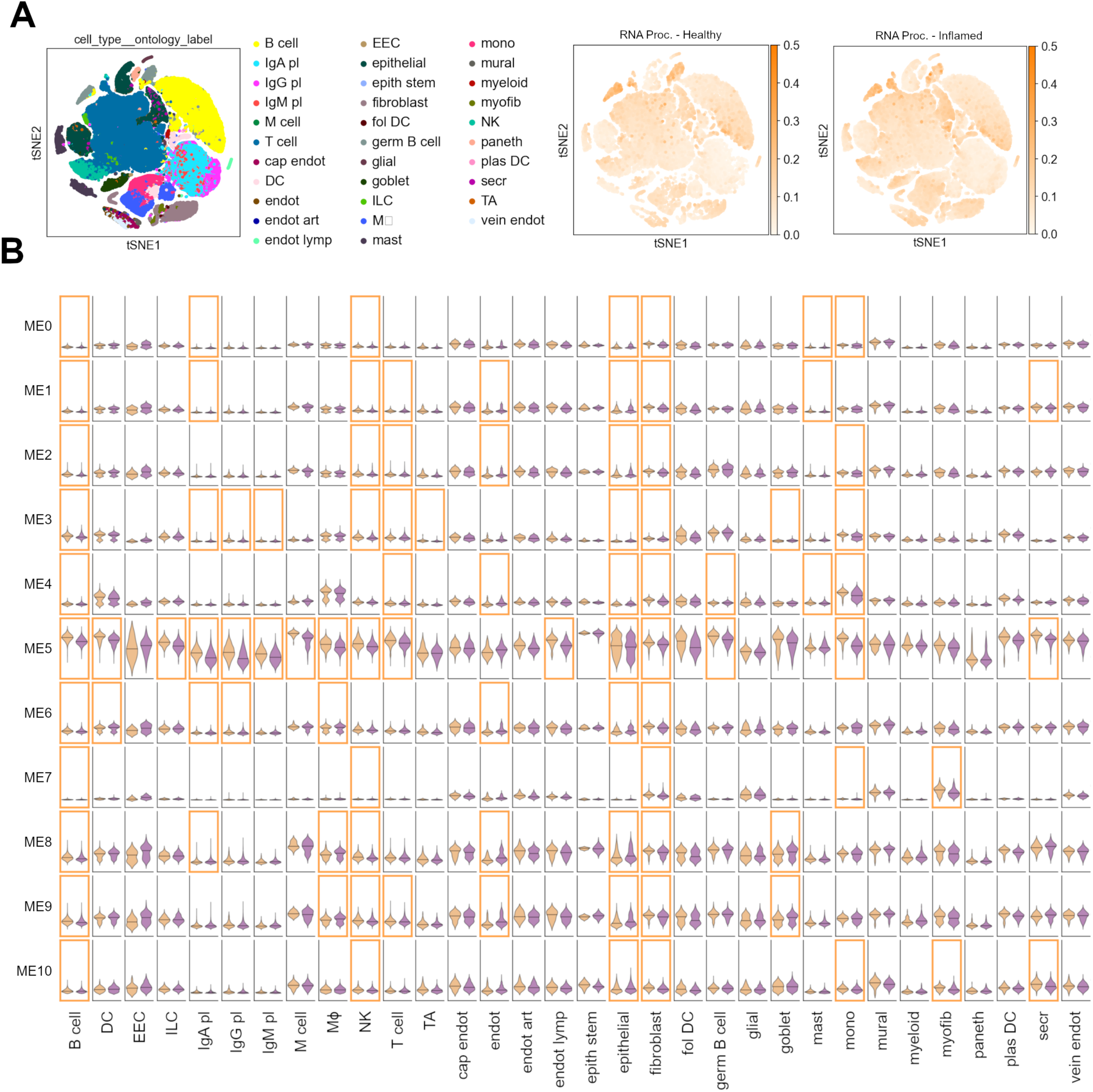

## Notes

https://www.ebi.ac.uk/biostudies/arrayexpress/

http://trr241.hosting.rrze.uni-erlangen.de/SEPIA/

https://www.ncbi.nlm.nih.gov/geo/

https://github.com/MiguelGonzalezAcera/SEPIA

### Competing Interest Statement

The authors have declared no competing interest.

### Summary of Updates

- Novel submission required editions. - Figure 2 is now in the Supplementary material. All its associated text as well. - Multiple sections in the text have been optimized.

## References

1 Neurath MF. Host-microbiota interactions in inflammatory bowel disease. Nat Rev Gastroenterol Hepatol 2020;17:76–7.

2 Neurath MF. Targeting cytokines in inflammatory bowel disease. Sci Transl Med 2022;14:eabq4473.

3 Friedrich M, Pohin M, Powrie F. Cytokine Networks in the Pathophysiology of Inflammatory Bowel Disease. Immunity 2019;50:992–1006.

4 Sazonovs A, Stevens CR, Venkataraman GR, Yuan K, Avila B, Abreu MT, et al. Large-scale sequencing identifies multiple genes and rare variants associated with Crohn’s disease susceptibility. Nat Genet 2022;54:1275–83.

5 de Lange KM, Moutsianas L, Lee JC, Lamb CA, Luo Y, Kennedy NA, et al. Genome-wide association study implicates immune activation of multiple integrin genes in inflammatory bowel disease. Nat Genet 2017;49:256–61.

6 Uhlig HH, Powrie F. Translating Immunology into Therapeutic Concepts for Inflammatory Bowel Disease. Annu Rev Immunol 2018;36:755–81.

7 Wirtz S, Popp V, Kindermann M, Gerlach K, Weigmann B, Fichtner-Feigl S, et al. Chemically induced mouse models of acute and chronic intestinal inflammation. Nat Protoc 2017;12:1295–309.

8 Gunther C, Martini E, Wittkopf N, Amann K, Weigmann B, Neumann H, et al. Caspase-8 regulates TNF-alpha-induced epithelial necroptosis and terminal ileitis. Nature 2011;477:335–9.

9 Ostanin DV, Pavlick KP, Bharwani S, D’Souza D, Furr KL, Brown CM, et al. T cell-induced inflammation of the small and large intestine in immunodeficient mice. Am J Physiol Gastrointest Liver Physiol 2006;290:G109–19.

10 Kontoyiannis D, Pasparakis M, Pizarro TT, Cominelli F, Kollias G. Impaired on/off regulation of TNF biosynthesis in mice lacking TNF AU-rich elements: implications for joint and gut-associated immunopathologies. Immunity 1999;10:387–98.

11 Roberts SJ, Smith AL, West AB, Wen L, Findly RC, Owen MJ, et al. T-cell alpha beta + and gamma delta + deficient mice display abnormal but distinct phenotypes toward a natural, widespread infection of the intestinal epithelium. Proc Natl Acad Sci U S A 1996;93:11774–9.

12 Fox JG, Ge Z, Whary MT, Erdman SE, Horwitz BH. Helicobacter hepaticus infection in mice: models for understanding lower bowel inflammation and cancer. Mucosal Immunol 2011;4:22–30.

13 Borenshtein D, McBee ME, Schauer DB. Utility of the Citrobacter rodentium infection model in laboratory mice. Curr Opin Gastroenterol 2008;24:32–7.

14 Liu CY, Girish N, Gomez ML, Dube PE, Washington MK, Simons BD, et al. Transitional Anal Cells Mediate Colonic Re-epithelialization in Colitis. Gastroenterology 2022;162:1975–89.

15 Sturm G, Finotello F, Petitprez F, Zhang JD, Baumbach J, Fridman WH, et al. Comprehensive evaluation of transcriptome-based cell-type quantification methods for immuno-oncology. Bioinformatics 2019;35:i436–i45.

16 Gu Z, Hubschmann D. simplifyEnrichment: A Bioconductor Package for Clustering and Visualizing Functional Enrichment Results. Genomics Proteomics Bioinformatics 2023;21:190–202.

17 Sanchez D, Batet M. Semantic similarity estimation in the biomedical domain: an ontology-based information-theoretic perspective. J Biomed Inform 2011;44:749–59.

18 Liberzon A, Birger C, Thorvaldsdottir H, Ghandi M, Mesirov JP, Tamayo P. The Molecular Signatures Database (MSigDB) hallmark gene set collection. Cell Syst 2015;1:417–25.

19 Ritchie ME, Phipson B, Wu D, Hu Y, Law CW, Shi W, et al. limma powers differential expression analyses for RNA-sequencing and microarray studies. Nucleic Acids Res 2015;43:e47.

20 Hanzelmann S, Castelo R, Guinney J. GSVA: gene set variation analysis for microarray and RNA-seq data. BMC Bioinformatics 2013;14:7.

21 Castanza AS, Recla JM, Eby D, Thorvaldsdottir H, Bult CJ, Mesirov JP. Extending support for mouse data in the Molecular Signatures Database (MSigDB). Nat Methods 2023;20:1619–20.

22 Subramanian A, Tamayo P, Mootha VK, Mukherjee S, Ebert BL, Gillette MA, et al. Gene set enrichment analysis: a knowledge-based approach for interpreting genome-wide expression profiles. Proc Natl Acad Sci U S A 2005;102:15545–50.

23 Langfelder P, Horvath S. WGCNA: an R package for weighted correlation network analysis. BMC Bioinformatics 2008;9:559.

24 Fernandez-Marcos PJ, Auwerx J, Schoonjans K. Emerging actions of the nuclear receptor LRH-1 in the gut. Biochim Biophys Acta 2011;1812:947–55.

25 Yi Q, Wang J, Song Y, Guo Z, Lei S, Yang X, et al. Ascl2 facilitates IL-10 production in Th17 cells to restrain their pathogenicity in inflammatory bowel disease. Biochem Biophys Res Commun 2019;510:435–41.

26 Miller KD, O’Connor S, Pniewski KA, Kannan T, Acosta R, Mirji G, et al. Acetate acts as a metabolic immunomodulator by bolstering T-cell effector function and potentiating antitumor immunity in breast cancer. Nat Cancer 2023;4:1491–507.

27 Yao Y, Cai X, Fei W, Ye Y, Zhao M, Zheng C. The role of short-chain fatty acids in immunity, inflammation and metabolism. Crit Rev Food Sci Nutr 2022;62:1–12.

28 Peluso AA, Kempf SJ, Verano-Braga T, Rodrigues-Ribeiro L, Johansen LE, Hansen MR, et al. Quantitative Phosphoproteomics of the Angiotensin AT(2)-Receptor Signalling Network Identifies HDAC1 (Histone-Deacetylase-1) and p53 as Mediators of Antiproliferation and Apoptosis. Hypertension 2022;79:2530–41.

29 Skarie JM, Link BA. The primary open-angle glaucoma gene WDR36 functions in ribosomal RNA processing and interacts with the p53 stress-response pathway. Hum Mol Genet 2008;17:2474–85.

30 Wilkins BJ, Lorent K, Matthews RP, Pack M. p53-mediated biliary defects caused by knockdown of cirh1a, the zebrafish homolog of the gene responsible for North American Indian Childhood Cirrhosis. PLoS One 2013;8:e77670.

31 Moudry P, Chroma K, Bursac S, Volarevic S, Bartek J. RNA-interference screen for p53 regulators unveils a role of WDR75 in ribosome biogenesis. Cell Death Differ 2022;29:687–96.

32 Kinchen J, Chen HH, Parikh K, Antanaviciute A, Jagielowicz M, Fawkner-Corbett D, et al. Structural Remodelling of the Human Colonic Mesenchyme in Inflammatory Bowel Disease. Cell 2018;175:372–86 e17.

33 Ogata H, Goto S, Sato K, Fujibuchi W, Bono H, Kanehisa M. KEGG: Kyoto Encyclopedia of Genes and Genomes. Nucleic Acids Res 1999;27:29–34.

34 Kanehisa M, Goto S. KEGG: kyoto encyclopedia of genes and genomes. Nucleic Acids Res 2000;28:27–30.

35 Li J, Simmons AJ, Chiron S, Ramirez-Solano MA, Tasneem N, Kaur H, et al. A Specialized Epithelial Cell Type Regulating Mucosal Immunity and Driving Human Crohn’s Disease. bioRxiv 2023.

36 Smillie CS, Biton M, Ordovas-Montanes J, Sullivan KM, Burgin G, Graham DB, et al. Intra-and Inter-cellular Rewiring of the Human Colon during Ulcerative Colitis. Cell 2019;178:714–30 e22.

37 Zheng HB, Doran BA, Kimler K, Yu A, Tkachev V, Niederlova V, et al. Concerted changes in the pediatric single-cell intestinal ecosystem before and after anti-TNF blockade. medRxiv 2023:2021.09.17.21263540.

38 VanDussen KL, Stojmirovic A, Li K, Liu TC, Kimes PK, Muegge BD, et al. Abnormal Small Intestinal Epithelial Microvilli in Patients With Crohn’s Disease. Gastroenterology 2018;155:815–28.

39 Quraishi MN, Acharjee A, Beggs AD, Horniblow R, Tselepis C, Gkoutos G, et al. A Pilot Integrative Analysis of Colonic Gene Expression, Gut Microbiota, and Immune Infiltration in Primary Sclerosing Cholangitis-Inflammatory Bowel Disease: Association of Disease With Bile Acid Pathways. J Crohns Colitis 2020;14:935–47.

40 Haberman Y, Karns R, Dexheimer PJ, Schirmer M, Somekh J, Jurickova I, et al. Ulcerative colitis mucosal transcriptomes reveal mitochondriopathy and personalized mechanisms underlying disease severity and treatment response. Nat Commun 2019;10:38.

41 Pelia R, Venkateswaran S, Matthews JD, Haberman Y, Cutler DJ, Hyams JS, et al. Profiling non-coding RNA levels with clinical classifiers in pediatric Crohn’s disease. BMC Med Genomics 2021;14:194.

42 Frede A, Czarnewski P, Monasterio G, Tripathi KP, Bejarano DA, Ramirez Flores RO, et al. B cell expansion hinders the stroma-epithelium regenerative cross talk during mucosal healing. Immunity 2022;55:2336–51 e12.

43 Haberman Y, Tickle TL, Dexheimer PJ, Kim MO, Tang D, Karns R, et al. Pediatric Crohn disease patients exhibit specific ileal transcriptome and microbiome signature. J Clin Invest 2014;124:3617–33.

44 van de Veen W, Globinska A, Jansen K, Straumann A, Kubo T, Verschoor D, et al. A novel proangiogenic B cell subset is increased in cancer and chronic inflammation. Sci Adv 2020;6:eaaz3559.

45 Artyomov MN, Van den Bossche J. Immunometabolism in the Single-Cell Era. Cell Metab 2020;32:710–25.

46 Barnhoorn MC, Hakuno SK, Bruckner RS, Rogler G, Hawinkels L, Scharl M. Stromal Cells in the Pathogenesis of Inflammatory Bowel Disease. J Crohns Colitis 2020;14:995–1009.

47 Brown SL, Riehl TE, Walker MR, Geske MJ, Doherty JM, Stenson WF, et al. Myd88-dependent positioning of Ptgs2-expressing stromal cells maintains colonic epithelial proliferation during injury. J Clin Invest 2007;117:258–69.

48 Miyoshi H, VanDussen KL, Malvin NP, Ryu SH, Wang Y, Sonnek NM, et al. Prostaglandin E2 promotes intestinal repair through an adaptive cellular response of the epithelium. EMBO J 2017;36:5–24.

49 Roulis M, Kaklamanos A, Schernthanner M, Bielecki P, Zhao J, Kaffe E, et al. Paracrine orchestration of intestinal tumorigenesis by a mesenchymal niche. Nature 2020;580:524–9.

