## Supplemental Information for "Integrated multi-model analysis of intestinal inflammation exposes key molecular features of preclinical and clinical IBD"

**Supplementary information**

**Supplementary Results**

**Transcriptomic correlations among models reveal overlapping intra- and inter-category regulation (relates to Fig. 2)**

Having identified how different inflammatory onsets influence gut cellular composition, we conducted transcriptome-wide analyses of similarities and differences. Transcriptome-transcriptome correlation among the colonic and ileal models revealed the proportion of intra- and inter-category regulatory overlap (Supplemental Fig S3A, B). Models with the greatest transcriptomic impact were AcDSS (barrier damage), cTNBS (immune modulation), and Hhepa (infectious) Supplemental Fig S3A, diagonal pie charts). Within barrier damage models, AcDSS and cDSS showed moderate correlation of determination (R^2^ = 0.65). Interestingly, cDSS and *Casp8*^ΔIEC^Col had the highest correlation within the barrier damage category (R^2^ = 0.67), suggesting similar regulatory changes induced by chronic DSS administration and persistent IEC necroptosis in *Casp8*^ΔIEC^Col (Supplemental Fig S3A). Some of the immune-driven colitis models also reached high correlation (OxC versus cTNBS, R^2^ = 0.79) (Supplemental Fig S3A). However, the highest intra-category correlation was within the infectious colitis category (R^2^ = 0.86), despite the differing requirements for IL-10 receptor inhibition in the Hhepa versus the Crode models. (Supplemental Fig. S3A).

Besides intra-category correlations, several high-level inter-category correlations were also identified (R2 > 0.7), notably between the barrier model *Casp8*ΔIECCol and the infection model Crode (R2 = 0.8) and the immune model TC (R2 = 0.7) (Supplemental Fig S3A). The infection model Crode showed some of the highest inter-category correlations against the immune models OxC and TC (R2 = 0.76 and 0.71, respectively) (Supplemental Fig S3A).

Interestingly, despite the size of the regulated transcriptome in the small intestinal models being fairly large (Supplemental Fig. S3B, diagonal pie charts), correlations remained low. The highest correlation was between *Casp8*^ΔIEC^Ile and *Tnf*^ΔARE^Ile (R^2^ = 0.52) (Supplemental Fig. S3B), indicating distinct and poorly concordant transcriptomic responses to inflammation triggers in the small intestine.

These analyses reveal contrasts in different model categories of gut inflammation. While anticipated similarities within each category were observed, significant cross-category similarities suggest common regulatory overlaps regardless of the nature of inflammatory onset.

**Supplementary Methods**

**Mice**

All mice on a C57BL/6J background were purchased from Charles River (Charles River GmbH) and Janvier Labs (Janvier Labs GmbH) and were housed under specific pathogen-free (SPF) conditions. Sample sizes for each mouse model have been empirically determined, the details of sample sizes are given in supplemental table 1. None of the animals were excluded in the analyses presented here, except for the WGCNA analysis where the TC experiment including controls were excluded to avoid data skewing due to confounder bias in the absence of T and B lymphocytes.

**Acute Dextran Sulfate Sodium (DSS)-induced colitis**

Five wild-type mice with a C57BL/6J background were subjected to 2% DSS with molecular weight ranging from 36,000 – 50,000 Da (MP Biomedicals) dissolved in drinking water for eight days. Evaluation of symptoms of inflammation and colitis parameters was performed as indicated in previous studies [1]. The mice were sacrificed on day nine to obtain inflamed colonic tissue for RNA sequencing.

**Chronic DSS-induced colitis**

C57BL/6J mice were subjected to three one-week (7 day) cycles of 2% DSS (M.W. range 36,000 – 50,000 Da, MP Biomedicals) interspersed by 2, two-week cycles of recovery. Mice were euthanized 10 days post-the last DSS cycle, and their colon was collected for RNA sequencing of the tissue.

**Oxazolone-induced colitis**

The induction of inflammation in the oxazolone model requires a preliminary sensitization step, whereby 3% (wt/vol) solution of 4-Ethoxymethylene-2-phenyl-2-oxazolin-5-one (oxazolone) (Sigma-Aldrich) in a 1:4 dilution with olive oil / acetone mixture is applied to the shaved skin of the animal, as previously described [2]. Seven days following the sensitization procedure, intrarectal enema administration of 150 µl of 0.5% oxazolone in 50% ethanol was conducted. Mice were sacrificed 24 hours later and the colonic tissues were dissected and snap frozen in liquid nitrogen using cryovials for RNA sequencing and fixed in 4% Histofix (Carl Roth GmbH) for histology.

**Acute TNBS-induced colitis**

Colitis was induced by the hapten TNBS (2,4,6-trinitrobenzenesulfonic acid, Merck) as previously described [2]. Mice were sensitized by epicutaneous application of 1% TNBS at a dilution of 1:4 in a mixture of oil and acetone (100 μl) on day 0, followed by intrarectal administration of 2% TNBS in 45% ethanol (100 μl) on day 6, followed by monitoring of body weight and mini-endoscopy. The mice were euthanized and colonic tissues were harvested and snap frozen in cryovials for RNA Sequencing and fixed in 4% Histofix (Karl Roth GmbH) for histology.

**Chronic TNBS-induced colitis**

For the induction of chronic colitis by TNBS, mice were sensitized as above [2] with 1% of TNBS in a mixture of oil and acetone (1:4). Subsequently, mice were administered with 150 µl of 0.5% TNBS via rectal enema on a 14-day schedule over a three-cycle period. The development of colitis was monitored over time by means of mini-endoscopy. At the end of the experiment, mice were euthanized and colonic tissues were harvested, snap frozen and fixed as above.

**T-cell transfer colitis**

For this model, a special control group of immune-deficient *Rag1*^-/-^ mice, coming from a C57BL/6J background is necessary. For the inflammation model, naïve CD4^+^ CD25^-^ T-cells were isolated from the spleen of wild-type C57BL/6J mice using MACS-based isolation (CD4^+^ T Cell Isolation Kit, mouse, Miltenyi Biotec) with additional CD25 negative selection (CD25 MicroBead Kit, mouse, Miltenyi Biotec). The obtained cells were controlled for their viability and purity via flow cytometry. One million cells were injected intraperitoneally in *Rag1*^-/-^ mice on a C57BL/6J background. The development of colitis was monitored over time via assessment of weight loss and mini-coloscopy. Colonic tissue for RNA sequencing was extracted from the sacrificed mice 3 weeks after the inoculation.

***Caspase8*^ΔIEC^ ileitis and colitis**

*Casp8*^ΔIEC^ mice were generated by crossing mice carrying a loxP-flanked caspase-8 allele (*Casp8*fl) mice to Villin-Cre mice, which were described earlier [3]. Cre-mediated recombination was genotyped by polymerase chain reaction on tail DNA. The development of ileitis and colitis has been previously characterized [3]. The onset of ileitis was observed to occur approximately at week 12, with the mice being sacrificed at approximately week 14. Mouse disease burden was assessed regularly according to the FELASA guidelines.

***Tnf*^ΔARE^ ileitis**

The *Tnf*^ΔARE^ mouse model is a widely used genetic model for studying Crohn's disease. *Tnf*^ΔARE^ mice are characterized by the deletion of the AU-rich element (ARE) in the tumor necrosis factor-alpha (TNF) gene, resulting in the constitutive overexpression of TNF. This leads to the development of chronic tissue inflammation, particularly affecting the terminal ileum. The *Tnf*^ΔARE^ mouse model closely resembles human Crohn's disease in terms of its histological features and the shared pathogenetic role of TNF. Mice heterozygous for the *Tnf*^ΔARE^ modification start to develop histological abnormalities at the age of 6 weeks [4]. For this study, 12-week-old heterozygous mice with fully established signs of ileal inflammation were used. The mouse line was a kind gift by Fabio Cominelli (Case Western Reserve University, USA).

***Eimeria vermiformis* infection**

Oocysts from *E. vermiformis* provided by Marc Veldhoen, (Instituto de Medicina Molecular, Faculdade de Medicina da Universidade de Lisboa, Portugal) were obtained and stored in 2.5% potassium bicromate, according to previously published protocols [5]. The oocysts were washed three times with deionized water, centrifuged at 1,800 g for 8 minutes, and floated at 1,100 g for 10 minutes. A final sterilization was performed using sodium hypochlorite. After three more washing cycles with deionized water, they were counted with the help of a Fuchs-Rosenthal chamber. One thousand of the cleaned oocysts diluted in water were used for infection of the mice by oral gavage. The mice were sacrificed nine days after the infection, and the ileal tissue was collected for RNA sequencing.

***Helicobacter hepaticus* infection**

*H. hepaticus* (DSMZ no.: 22909) was revived in glycerol stocks and cultured on blood agar plates under microaerobic conditions at 37°C. Microscopic examination confirmed the bacterial quality and purity before the cultures were transferred to tryptone soya broth supplemented with 10% fetal calf serum, 10 μg/mL vancomycin, 5 μg/mL trimethoprim, and 2.5 IU/mL polymyxin B. Cultivation proceeded under microaerophilic conditions at 37°C, with shaking at 180 rpm. Mice were orally inoculated with 1 × 10^8^ colony-forming units (c.f.u.) of *H. hepaticus* on two consecutive days using a 22 G curved blunted needle. Concurrently, mice received intraperitoneal injections (i.p.) of 1 mg of anti-IL-10R antibody (clone 1B1.2, BioXcell) on day 0 and at weekly intervals thereafter [6, 7, 8]. On day 14 post-inoculation, mice were euthanized, and colon samples were harvested for subsequent analyses. Sections from colon were preserved in RNA Later (Qiagen) for gene expression studies. Histological evaluation of colitis was carried out in accordance with the methodology previously described [9].

***Citrobacter rodentium* infection**

For this model we used the ICC169 strain of *C. rodentium*, as referenced in previous studies [10]. The organism was cultivated in a sterilized LB medium at a temperature of 37°C under constant aeration and shaking conditions. The culture was supplemented with erythromycin. For the inoculation of the bacteria, mice were subjected to eight hours of fasting, after which ~10^9^ colony-forming units (CFU) of *C. rodentium,* diluted in sterile phosphate-buffered saline (PBS), were introduced into the animal via oral gavage. Eight days later, the mice were sacrificed and their colons were collected for RNA sequencing.

Experimental protocols were approved by the Institutional Animal Care and Use Committee of the Regierung von Unterfranken and the Landesamt für Gesundheit und Soziales – Berlin, and in accordance with the UK Scientific Procedures Act of 1986.

**Human cohort data**

Publicly available IBD patient cohorts that were used in this study include the treatment naïve cohorts GSE109142 (PROTECT), GSE117993 (RISK-UC) [11], GSE57945 (RISK - CD) [12], cohort of adult IBD with concurrent PSC E-MTAB 7915 (PSC-UC) [13], and the adult Crohn’s disease patient cohort E-MTAB 5783 (WashU) [14]. For the in-house cohort (IBDome), samples were obtained from non-IBD and IBD patients after written informed consent was obtained from the participants at the Gastroenterology Department, of Charite – Universitätsmedizin Berlin as well as the Universitätsklinikum Erlangen. Biopsies were taken during endoscopy or removed from resected tissue and stored in 10 mL RNA protect Tissue Reagent (Qiagen) overnight at 4°C. For long-term storage, the reagent was removed and biopsies kept at -80°C. For RNA isolation, the RNeasy Kit (Qiagen) was used according to the manufacturer´s instructions employing the TissueLyser LT (Qiagen) and 5 mm balls. Concentration and purity of the isolated RNA was determined by the ratios of absorbance (A260:280 and A260:230) using NanoDrop 1000 (Thermo Fisher Scientific). The RNA Integrity Number (RIN) was determined using the Agilent RNA ScreenTape System (Agilent Technologies) according to manufacturer’s instructions employing the Agilent 2200 TapeStation. The RNA Clean & Concentrator™-25 (Zymo Research) was used for RNA samples with a RIN < 1.8 according to the manufacturer´s instructions. RNA samples were shipped on dry ice to Quantitative Biology Center (QBiC) of Eberhard Karls Universität Tübingen for sequencing. All data from the patients were pseudonymized.

**Histological and immunohistological analyses**

For the hematoxylin and eosin (H&E) stainings, we prepared sections of 5-10 μm from tissues fixed in formalin and embedded in paraffin, which were then subjected to standard H&E staining protocol. The immunostainings were performed on paraffin sections of the tissues, which were cut, deparaffinized, hydrated, and treated with a Tris-EDTA-based antigen retrieval solution. To prevent non-specific binding, a commercial blocking reagent (Immunoblock 1X, Carl Roth GmbH) was utilized. The samples were then incubated overnight at 4°C in the dark with the primary antibody for each immunostaining (table below). We washed the samples in TBS and then we incubated them with either the secondary antibody or streptavidin conjugates (DyLight, Invitrogen) for 1 h at 4°C. Samples were then counterstained with Hoechst 33342 (Thermo Fisher Scientific) for the cellular nuclei and cover-slipped in fluorescence mounting medium. A Leica TCS SP5 confocal microscope (Leica Microsystems) was employed for the acquisition of the immunofluorescence images using the necessary settings.

| **Antibody** | **Type** | **Dilution** | **Species** | **Catalogue #** | **Company** |
| --- | --- | --- | --- | --- | --- |
| *Primary antibody* | | | | | |
| Anti-F4/80 | Unconjugated | 1:200 | Rabbit | 7007S | Cell Signaling |
| Anti-CD3 | Unconjugated | 1:100 | Rat | 100238 | Biolegend |
| Anti-KI67 | Alex Fluo555 | 1:100 | Rat | 14-5698-82 | abcam |
| Anti-MUC2 | Unconjugated | 1:200 | Rabbit | NBP1-31231 | Novus B |
| Anti-S100B | Unconjugated | Ready to use | Rabbit | Dako | Biolegend |
| Anti-TUBB3 | Unconjugated | 1:100 | Mouse | 801212 | Biolegend |
| Anti-COX4 |  | 1:2400 | Rabbit | 4844 | Cell Signaling |
| Anti-TOMM20 |  | 1:200 | Rabbit | HPA011562 | Sigma |
| *Secondary antibody* | | | | | |
| Anti-Rabbit | Biotinylated | 1:500 | Goat | 111-065-144 | Jackson |
| Anti-Rat | Biotinylated | 1:500 | Goat | 554014 | BD Biosciences |

**RNA extraction**

The RNA obtained from the collected tissue samples was extracted using multiple extraction kits, according to the manufacturer’s instructions, using the peqGold total tissue RNA kit (peqlab GmbH) and the MicroSpin total RNA kit (VWR International GmbH). The obtained RNA was controlled for contamination and degradation with the NanoPhotometer® spectrophotometer (IMPLEN, CA, USA), the Nanodrop (thermofischer), the Qbit (Thermo), and the RNA Nano 6000 Assay Kit of the Bioanalyzer 2100 system (Agilent Technologies, CA, USA).

**Library Preparation and mRNA Sequencing**

We aliquoted 1 μg of RNA per sample to prepare the libraries for sequencing. We used the NEBNext® UltraTM RNA Library Prep Kit for Illumina® (NEB, USA) according to the manufacturer's instructions, with indexing added to the molecules from each sample. From the total RNA of each sample, mRNA was purified using poly-T oligo-attached magnetic beads. The molecules were fragmented with divalent cations at high temperatures while diluted in NEBNext First Strand Synthesis Reaction Buffer (5X). We used a random hexamer primer and M-MuLV reverse transcriptase (RNase H-) to synthesize the first strand of cDNA, and DNA polymerase I was followed to synthesize the second strand. Excess RNA was cleared using RNase H. The overhang of the molecules was blunted through the action of the exonuclease/polymerase. The 3’ ends of the DNA fragments were adenylated and a NEBNext adaptor with a hairpin loop structure was ligated to them to prepare for hybridization. The AMPure XP system (Beckman Coulter, Beverly, USA) was used to purify the fragments of 150-200 bp in length. To prepare the fragments for PCR, we used 3 μl of USER Enzyme (NEB, USA) at 37°C for 15 minutes and 5 minutes at 95°C. We performed the PCR with Phusion High-Fidelity DNA polymerase, Universal PCR primers and Index (X) Primer. The products of the PCT were purified again with the AMPure XP system and the resulting library was controlled for quality using the Agilent Bioanalyzer 2100 system.

We used a cBot Cluster Generation System using PE Cluster Kit cBot-HS (Illumina) for the clustering of the indexed samples according to the manufacturer's instructions. The prepared libraries were sequenced using an Illumina platform, generating paired-end reads. Initial trimming of the adapter and poly-N sequences, as well as demultiplexing of the reads was done by the Illumina software included in the platform. Further processing of the raw data (raw reads) was done using in-house scripts and publicly available software. Further analysis was performed in R software for statistical programming (4.4.0). Quality control of the fastq files was performed using FastQC (v0.12.1), calculating Q20, Q30 and GC content. We downloaded the current mouse reference genome (GRCm39) and its annotation from NCBI, UCSC and Ensembl. The genome was indexed and used for mapping the paired-end reads using STAR mapping software (2.7.10b) followed by sequence alignments using picard (3.1.1) and samtools (1.20) and the analysis of counts per gene using featureCounts (v2.0.6).

**Analysis of ontology based semantic-similarity networks**

To uncover the functional attributes and the relatedness among these shared genes, we performed an analysis for ontology-based semantic similarity on these genes (24-26). This method yields a network of sub-clusters where the nodes are ontologies and the edges are the kappa similarity coefficients. These analyses were performed as described by Sanchez *et al*. in their original article [15]. Clustering was performed on all terms reaching significance and with a kappa similarity of above 0.3. Cytoscape (3.10.2) was used for the visualization of the similarity network.

**Deconvolution of gene expression**

Processing of publicly available scRNA-Seq datasets was performed using scanpy (1.10.1) in python (3.10.12), using the provided data and annotation. The DWLS package (0.1.0) was used for the cell proportion deconvolution analysis. GSVA (1.52.2) was employed for the hallmark pathway analysis, with the genes extracted from GSEA database. Differential analyses of resulting values were performed using limma (3.60.2).

**Analysis of differential gene expression**

Differential expression analysis was performed using the DESeq2 R package (1.44.0), filtering out genes with less than 15 counts overall. Over representation analyses (ORA) using the Gene Ontology and KEGG enrichment databases was performed using the ClusterProfiler R package (4.12.0).

**Weighted gene correlation network analysis**

We used the WGCNA package (1.72-5) [16] on the raw counts to obtain co-expression modules across samples, segregated by tissue. Default parameters and a soft threshold was chosen as maximum for every analyses. A post-clustering comparison of these module eigenvalues using limma-based linear modeling was performed as shown previously [17], yielding the differential changes per module across treatment groups, which we represented in our study as Δ eigenvalues. An ORA was performed using the database Gene Ontology to observe processes represented in each module using the ClusterProfiler R package (4.12.0).

**Analysis of publicly available scRNA-Seq data**

Orthogonal analysis of human IBD scRNA-Seq datasets (UC accession No. SCP259 and CD accession No SCP1423) to identify cell clusters expressing the transposed mouse module data was performed using scanpy (1.10.1) in python (3.10.12).

**Generation of figure plots**

The commercial software Graphpad Prism 9 and in-house python (3.11.0rc1) and R (4.4.0) scripts were used to generate the graphical displays. R package ComplexHeatmap (2.20.0) was used for the generation of heatmaps, Ridge plots were generated using the ggridges package (0.5.6). Other plots in R were generated using the ggplot2 package (3.5.1). Python package pyvenn was used for the generation of the venn diagrams. Matplotlib (3.9.0) and Seaborn (0.13.2) were used for the figure plots generated using python (3.11.0rc1). Cytoscape (3.10.2) was used for network data visualization.

Ancillary packages and dependencies used in data processing, transformation and management:

| **Package** | **Version** |
| --- | --- |
| *Python packages* | |
| Pandas | 2.2.2 |
| numpy | 1.26.4 |
| scipy | 1.13.1 |
| snakemake | 7.32.4 |
| flask | 3.0.3 |
| mysql.connector | 8.4.0 |
| reportlab | 4.2.0 |
| adjustText | 1.1.1 |
| *R packages* | |
| dendextend | 1.17.1 |
| cluster | 2.1.6 |
| gplots | 3.1.3.1 |
| circlize | 0.4.16 |
| gsubfn | 0.7 |
| GOstats | 2.70.0 |
| enrichplot | 1.24.0 |
| pathview | 1.44.0 |
| simplifyEnrichment | 1.14.0 |
| magrittr | 2.0.3 |

**Supplementary Figure legends**

**Supplementary Figure 1. Preclinical evaluation of mouse models.** **A)** Representative H&E histology images of the ileitis mouse models and healthy controls along with the indicated pre- and post-euthanasia endpoints for disease establishment. **B)** Representative images from pre-euthanasia screening of inflammation (top row: colonoscopy and IVIS Spectrum *in vivo* imaging), and post-euthanasia histological assessments (bottom row) from the colitis mouse models from each model category. **C)** Schematic illustration of the processing and analysis of the mouse-mouse and the mouse-human datasets. **D)** Principal component analyses (PCA) of the samples in the model databank (grey ellipse = colonic samples, yellow ellipse = ileal samples), few individual models have been highlighted.

**Supplementary Figure 2. Tissue composition across mouse models.** **A)** Bubble plot showing the expression of selected cell type markers across the ileitis mouse models. **B)** Ridge plots of the selected cell type markers across the ileitis mouse models.

**Supplementary Figure 3. Correlations between mouse models of A) colitis and B) ileitis.** Pie charts on the diagonal represent the number of genes that are significantly up- (orange) and down- (violet) regulated in each model and the legend with the empty bubbles indicates the number of genes. Scatter plots in the bottom left half show the linear regression between two given models, comparing the log2-fold change of common genes. The colour intensity of individual dots correlates directly with the slope of the regression line. Dot plots on the top right half represent the R^2^ value of the respective linear regression. Triangular outlines demarcate the comparisons within a given model category, namely barrier damage, immune modulation, or infection.

**Supplementary Figure 4: Exclusive processes observed in mouse models.** **A-B)** Exclusively up- A) and down- B) regulated genes from the colitis mouse models. The top three enriched Gene Ontologies are shown for each model. **C-D)** Exclusively up- C) and down- D) regulated genes from ileitis mouse models. The top three enriched Gene Ontologies are shown for each model.

**Supplementary Figure 5. Average expression of WGCNA modules across different single-cell clusters from UC samples**. **A-C)** Violin plots of average expression of the genes from each WGCNA module identified from the colitis models among the epithelial A), immune B), and stromal C) compartments, segregated by annotated cell type and patient health. Orange outlines indicate significant differences in expression in a two-tailed Mann-Whitney U test.

**Supplementary Figure 6. Translation of WGCNA modules and features from mouse ileitis models onto single-cell RNAseq from CD patients.** **A)** t-SNE of single cell RNAseq PREDICT experiment (left) and average expression of genes from the ontology ‘RNA processing’ included in mouse ileitis module ME2, segregated by inflammation status. **B)** Violin plots of average expression of the genes identified from each mouse ileitis WGCNA module segregated patient health and annotated by cell type. Orange outlines indicate significant differences in expression in a a two-tailed Mann-Whitney U test.

**Supplementary references**

1	Becker C, Fantini MC, Neurath MF. High resolution colonoscopy in live mice. Nat Protoc 2006;**1**:2900-4.

2	Wirtz S, Popp V, Kindermann M, Gerlach K, Weigmann B, Fichtner-Feigl S*, et al.* Chemically induced mouse models of acute and chronic intestinal inflammation. Nat Protoc 2017;**12**:1295-309.

3	Gunther C, Martini E, Wittkopf N, Amann K, Weigmann B, Neumann H*, et al.* Caspase-8 regulates TNF-alpha-induced epithelial necroptosis and terminal ileitis. Nature 2011;**477**:335-9.

4	Kontoyiannis D, Pasparakis M, Pizarro TT, Cominelli F, Kollias G. Impaired on/off regulation of TNF biosynthesis in mice lacking TNF AU-rich elements: implications for joint and gut-associated immunopathologies. Immunity 1999;**10**:387-98.

5	Figueiredo-Campos P, Ferreira C, Blankenhaus B, Veldhoen M. Eimeria vermiformis Infection Model of Murine Small Intestine. Bio Protoc 2018;**8**.

6	Fox JG, Ge Z, Whary MT, Erdman SE, Horwitz BH. Helicobacter hepaticus infection in mice: models for understanding lower bowel inflammation and cancer. Mucosal Immunol 2011;**4**:22-30.

7	Kullberg MC, Rothfuchs AG, Jankovic D, Caspar P, Wynn TA, Gorelick PL*, et al.* Helicobacter hepaticus-induced colitis in interleukin-10-deficient mice: cytokine requirements for the induction and maintenance of intestinal inflammation. Infect Immun 2001;**69**:4232-41.

8	West NR, Hegazy AN, Owens BMJ, Bullers SJ, Linggi B, Buonocore S*, et al.* Oncostatin M drives intestinal inflammation and predicts response to tumor necrosis factor-neutralizing therapy in patients with inflammatory bowel disease. Nat Med 2017;**23**:579-89.

9	Izcue A, Hue S, Buonocore S, Arancibia-Carcamo CV, Ahern PP, Iwakura Y*, et al.* Interleukin-23 restrains regulatory T cell activity to drive T cell-dependent colitis. Immunity 2008;**28**:559-70.

10	Riedel CU, Casey PG, Mulcahy H, O'Gara F, Gahan CG, Hill C. Construction of p16Slux, a novel vector for improved bioluminescent labeling of gram-negative bacteria. Appl Environ Microbiol 2007;**73**:7092-5.

11	Haberman Y, Karns R, Dexheimer PJ, Schirmer M, Somekh J, Jurickova I*, et al.* Ulcerative colitis mucosal transcriptomes reveal mitochondriopathy and personalized mechanisms underlying disease severity and treatment response. Nat Commun 2019;**10**:38.

12	Haberman Y, Tickle TL, Dexheimer PJ, Kim MO, Tang D, Karns R*, et al.* Pediatric Crohn disease patients exhibit specific ileal transcriptome and microbiome signature. J Clin Invest 2014;**124**:3617-33.

13	Quraishi MN, Acharjee A, Beggs AD, Horniblow R, Tselepis C, Gkoutos G*, et al.* A Pilot Integrative Analysis of Colonic Gene Expression, Gut Microbiota, and Immune Infiltration in Primary Sclerosing Cholangitis-Inflammatory Bowel Disease: Association of Disease With Bile Acid Pathways. J Crohns Colitis 2020;**14**:935-47.

14	VanDussen KL, Stojmirovic A, Li K, Liu TC, Kimes PK, Muegge BD*, et al.* Abnormal Small Intestinal Epithelial Microvilli in Patients With Crohn's Disease. Gastroenterology 2018;**155**:815-28.

15	Sanchez D, Batet M. Semantic similarity estimation in the biomedical domain: an ontology-based information-theoretic perspective. J Biomed Inform 2011;**44**:749-59.

16	Langfelder P, Horvath S. WGCNA: an R package for weighted correlation network analysis. BMC Bioinformatics 2008;**9**:559.

17	Liu Y. CWGCNA: an R package to perform causal inference from the WGCNA framework. NAR Genom Bioinform 2024;**6**:lqae042.
